# Expression of the β*hpmeh* gene in transgenic events of the potato variety Desiree increases resistance to bacterial wilt caused by *Ralstonia solanacearum*

**DOI:** 10.64898/2026.01.14.699457

**Authors:** Myriam Izarra, Liliam Gutarra, Elizabeth Fernández, Eva Huamán, Marc Ghislain, Jan Kreuze

## Abstract

Bacterial wilt, caused by *the Ralstonia solanacearum* species complex (RSSC), severely affects many important crops and significantly limits potato production worldwide. RSSC bacteria can regulate the expression of their virulence factors, including extracellular polysaccharides and endoglucanases, through quorum sensing, which enables bacteria to sense their population density through signaling molecules and collectively switch on virulence factors when a threshold density is reached. 3-hydroxy palmitic acid methyl ester is the main quorum-sensing molecule in RSSC that can be hydrolyzed by β-hydroxypalmytate methyl ester hydrolase (βHPMEH). In this study, we evaluated the ability of β*hpmeh* transgenic potato (Desiree) plants to reduce bacterial wilt symptoms after artificial inoculation with two virulent RSSC strains under controlled conditions and related this to the expression of the β*hpmeh* gene. For each transgenic event, we analyzed the phenotypic response (wilt incidence, latent infection, and area under the disease progress curve). Real-time quantitative PCR was performed to determine the relative expression levels of β*hpmeh* in transgenic events. Several transgenic events were identified with reduced susceptibility to bacterial wilt compared to Desiree and a resistance level similar to or higher than that of potato variety Cruza 148, the most resistant variety available, which was positively correlated with β*hpmeh* expression.

## Introduction

The *Ralstonia solanacearum* species complex (RSSC; E.F. Smith) comprises a group of soil-borne bacteria responsible for bacteria wilt, a destructive disease affecting numerous economically important crops, including potato, banana, cucurbits, eggplant, Eucalyptus, ginger, groundnut, mulberry, tobacco, tomato, and several ornamental plants. This pathogen poses a significant challenge to agricultural production across the globe due to its extensive host range and worldwide presence (Mansfield *et al*., 2012; Charkowski *et al*., 2020).

RSSC strains have historically been classified into five races and six biovars based on host range and metabolic characteristics (Denny & Hayward, 2001; Fegan & Prior, 2005; Ghorai *et al*., 2022). Advances in molecular analysis have further refined this framework, grouping strains into four major phylotypes (I-IV) according to sequence variation in conserved genes such as *egl*, *mutS*, *hrpB*, and ITS (Fegan & Prior, 2005). These phylotypes are broadly associated with their geographical origins, including Asia (phylotype I), the Americas (phylotype II, comprising IIA and IIB), Africa (phylotypes III) and Indonesia (phylotype IV)(Fegan & Prior, 2005; Paudel *et al*., 2020). More recently, taxonomic revisions have proposed dividing the complex into three species: *R. pseudosolanacearum* (phylotypes I and III), *R. solanacearum* (phylotype II), and *R. syzygii* (phylotype IV) (Safni *et al*., 2014). Within each phylotype, additional diversity is recognized at the sequevar level based on the *egl* gene sequences (Fegan & Prior, 2005; Chesneau *et al*., 2018; Ghorai *et al*., 2022). In potato, bacterial wilt (BW) is mainly associated with strains belonging to phylotype IIB, particularly sequevars 1 and 2, historically referred to as race 3/biovar 2A (Hayward, 1994).

In tropical potato production systems, BW, together with viral diseases, represents a major constraint, second only to late blight. Although several moderately resistant cultivars and wild potato species exhibiting quantitative resistance have been identified, complete resistance to tuber infection has not yet been achieved (Ferreira *et al*., 2017). Infected plants typically exhibit symptoms such as wilting, stunting, and leaf yellowing (Champoiseau *et al*., 2009).

In potato, *Ralstonia solanacearum* typically gains entry through wounds or natural openings in the roots, which may arise from soil friction, nematode activity, or the emergence of lateral roots (CABI, 2023). After entry, the bacterium colonizes the root cortex and proliferates within intercellular spaces. It then deploys a Type III secretion system (T3SS) to translocate effector proteins into host cells, thereby suppressing plant defense responses and enabling invasion of the xylem (Genin, 2010). Activation of the T3SS is thought to be triggered by plant-derived signals, including components of the cell wall, and is modulated by environmental conditions such as soil composition, temperature, and moisture (Peeters *et al*., 2013). In addition, certain compounds, such as oleanolic acid, have been reported to enhance T3SS activity, further promoting bacterial colonization (Wu *et al*., 2015).

Once inside the vascular system, *R. solanacearum* proliferates within the xylem vessels, where it produces major virulence determinants, including exopolysaccharides (EPS) and a variety of extracellular cell wall-degrading enzymes (CWDE) (Clough *et al*., 1997b; Flavier *et al*., 1997a,b; Schell, 2000). The accumulation of EPS contributes to the blockage of water transport, leading to wilting symptoms, while CWDE enhances virulence by facilitating invasion and vascular colonization (Saile *et al*., 1997). The production of these factors is controlled by a complex regulatory network that responds to environmental conditions and coordinates the expression of pathogenicity-related genes (Valls *et al*., 2006). This regulation is mediated by a quorum sensing system, in which gene expression is modulated according to cell densities (Clough *et al*., 1997b; Flavier *et al*., 1997a). As bacterial populations increase, a transition known as phenotype conversion occurs, shifting the bacterium from a motile state, associated with early infection and environmental survival, to a sessile, EPS-producing state adapted to vascular colonization and systemic spread (Denny & Hayward, 2001; Hikichi *et al*., 2007).

The phenotype conversion (*Phc*) regulon constitutes the central regulatory network controlling virulence gene expression in *R.solanacearum* (Clough *et al*., 1997a). A key component of this system is 3-hydroxypalmitic acid methyl ester (3-OH PAME), which is synthesized by the enzyme encoded by *phcB*, an S-adenosyl methionine-dependent methyltransferase that converts 3-hydroxypalmitic acid (3-OH PA) into its methyl ester form. This molecule functions as a quorum-sensing signal (quormone) that modulates the activity of the global regulator PhcA at the post-transcriptional level (Clough *et al*., 1997a). PhcA, a LysR-type transcriptional regulator, controls the production of EPS and other virulence factors and is associated with changes in bacterial motility (Fig. 1). When the concentration of 3-OH PAME exceeds approximately 5 nmol·L^-1^, PhcA becomes activated, leading to increased production of EPS and CWDE. Mutants in *phcA* exhibit reduced levels of EPS and extracellular proteins (EXPs), and are nearly avirulent (Brumbley *et al*., 1993). Similarly, *phcB* mutants show strongly reduced virulence, which can be restored by the addition of exogenous 3-OH PAME (Flavier *et al*., 1997a; Hikichi *et al*., 2017). These findings highlight the essential role of 3-OH PAME in regulating virulence in *R. solanacearum*.

**Figure 1.**
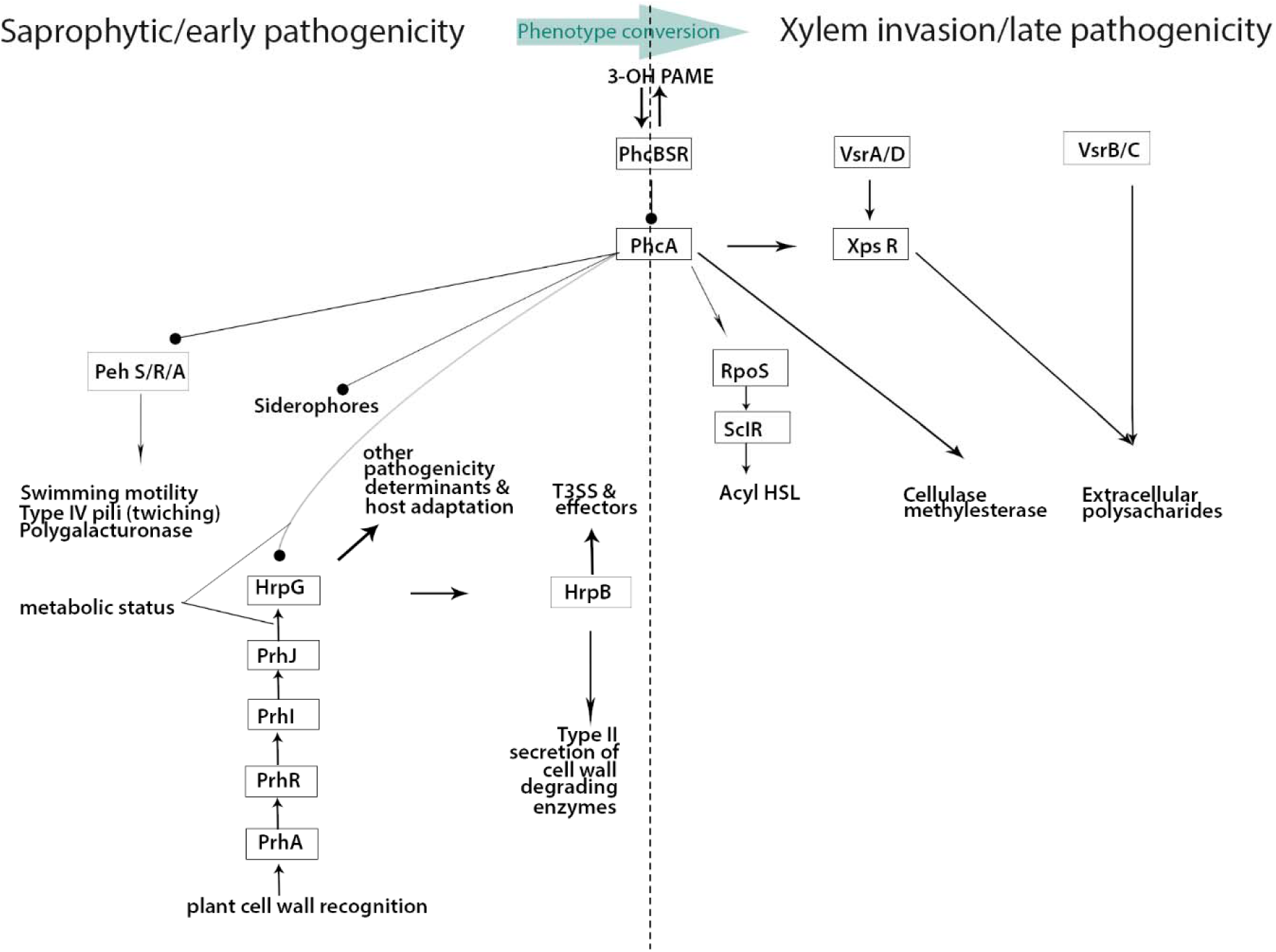
Simplified schematic representation of virulence regulation in *R. solanacearum*. PhcA is positioned at the center of the regulatory network and controls phenotypic conversion through the quorum-sensing signal 3-OH PAME. Arrows indicate activation, whereas lines ending in circles denote inhibition. T3SS: Type III secretion system.

Efforts to disrupt quorum sensing have been investigated as an alternative approach to reduce bacterial virulence. In many plant pathogenic bacteria, systems for cell-to-cell communication regulate the expression of genes involved in host colonization and disease progression (Whitehead *et al*., 2001). These systems rely on diffusible signaling molecules of diverse chemical nature, including N-acyl homoserine lactones (AHLs), fatty acid esters, and peptides. Disruption of these signaling pathways, referred to as quorum quenching, has been shown to attenuate virulence. For example, expression of AHL-lactonase enzymes, which inactivate AHL molecules by hydrolyzing their lactone bond, reduces the production of cell wall-degrading enzymes and diminishes pathogenicity in bacteria such as *Erwinia carotovora* (Dong *et al*., 2000). Furthermore, transgenic plants expressing AHL-lactonase exhibit enhanced resistance to bacterial infection (Dong *et al*., 2001). This quorum-quenching approach targets quorum-sensing systems by degrading signaling molecules, thereby preventing the coordinated expression of virulence factors and reducing pathogenicity (Zhang, 2003).

Nearly two decades ago, an enzyme, β-hydroxypalmitate methyl ester hydrolase (βHPMEH), was isolated from a soil bacterium and shown to exhibit high hydrolytic activity toward the ester bond of 3-OH PAME, thereby inactivating this signaling molecule and reducing the virulence of *R. solanacearum*. Crude enzyme extracts from recombinant *E. coli* were found to inhibit *in vitro* EPS production, suggesting that βHPMEH interferes with the activation of virulence-associated genes (Shinohara *et al*., 2007).

In this study, we developed transgenic potatoes expressing βHPMEH and evaluated their resistance to *Ralstonia solanacearum* under controlled greenhouse conditions. Two modes of expression were used: constitutive and tissue-specific. The 35S promoter of the cauliflower mosaic virus (CaMV) was used to drive strong constitutive expression, as documented in many transgenic plants (Amack & Antunes, 2020). For tissue-specific expression, the GRP1.8 promoter, which directs xylem-specific expression of a glycine-rich protein (GRP), was selected. This promoter, originally isolated from bean and characterized in tobacco, directs expression in the xylem parenchyma of young roots (Keller & Baumgartner, 1991). GRPs are structural cell wall proteins that contain an apoplastic secretion signal, enabling their secretion into the xylem and subsequent movement with the xylem sap (Lv *et al*., 2009). Tissue-specific expression of a quorum-quenching system is desirable in this strategy to minimize pleiotropic effects on plant metabolism while targeting key infection sites. Resistance to *R. solanacearum* was assessed through phenotypic and genotypic analyses.

## Materials and Methods

### Constructs and generation of transgenic events

The gene constructs used in this study (pPAMEH and pGRP1.8-PAMEH) have been described previously (Fernández *et al*., 2015). They comprise the plant codon-optimized βHPMEH gene, preceded by the GRP signal peptide for extracellular targeting to the apoplast and a viral translational enhancer signal, under the control of the constitutive 35S or xylem-specific GRP1.8 promoters. Potato transformation was performed according to the *Agrobacterium tumefaciens* organogenesis protocol described by Medina-Bolivar et al. (2003) with minor modifications for the variety Desiree. Pathogen-free plants of the potato variety Desiree (accession number CIP800048) were obtained from the potato germplasm collection of the International Potato Center (CIP). Plantlets were cultured in liquid propagation medium (4.3 g·L^-1^ Murashige and Skoog (MS) salts, 0.4 mg·L^-1^ thiamine, 2 m g·L^-1^ glycine, 0.5 mg·L^-1^ nicotinic acid, 0.5 mg·L^-1^ pyridoxine, 0.1 mg·L^-1^ gibberellic acid, 2% sucrose, and pH 5.6) during 4 weeks in a growth room at 14–16°C, 70% relative humidity, and 16 h photoperiod (2,000 lx). In parallel, *A. tumefaciens* carrying the plasmid pPAMEH or pGRP1.8-PAMEH were cultivated on Luria–Bertani (LB; 1% bacto-tryptone, 0.5% yeast extract, 1% NaCl, pH 7.5, and 2.5 g·L^-1^ of agar) supplemented with 100 mg·L^-1^ kanamycin at 28°C for 48 h. A single bacterial colony was used to inoculate 3 mL of LB liquid medium containing 100 mg·L^-1^ kanamycin and 100 mg·L^-1^ rifampicin, incubated at 28°C for 20 h in a water bath shaker (250 rpm). Leaves with the petiole (1.5–2 cm) were cut from the top third of the *in vitro* plantlets and co-cultivated with *A. tumefaciens* at a final concentration of 5.9 × 10^7^ cells·mL^-1^ (bacterial concentration was estimated from a standard curve by absorbance at 620 nm) in 25 ml of co-culture medium (4.6 g·L^-1^ of MS salts, 2% sucrose, and 50 µM acetosyringone) at 22°C for 24 h in the dark. The explants were then taken out from the liquid medium and briefly dried on sterile filter paper before transferring them to semi-solid regeneration medium (4.6 g·L^-1^ MS salts, 0.02 mg·L^-1^ gibberellic acid, 0.02 mg·L^-1^ naphthalene acetic acid, 2 mg·L^-1^ zeatin riboside, 2% sucrose, pH 5.6, and 4 g·L^-1^ of agar) without antibiotic for one day. Subsequently, the explants were transferred (ten per Petri dish) to a semi-solid regeneration medium containing 100 mg·L^-1^ kanamycin and 200 mg·L^-1^ carbenicillin and were transferred onto fresh medium every 2 weeks. After one month, regenerants started to appear from *Agrobacterium-infected* leaf explants, and 6-week-old putative transgenic regenerants (only one regenerant per explant) were screened for *nptII* (Km-F/R), *pameh 241* (F/R), *35S-pameh* (F/R), and *GRP1.8-pameh* (F/R) genes as described in Fernández et al. (2015). The regenerants were propagated in test tubes containing semi-solid regeneration medium at 15–18°C under artificial light (16 h light/ 8 h dark). Leaves from the putative transgenic regenerants were confirmed to be transgenic on a highly selective medium: leaves were laid on kanamycin selective callus-inducing medium (4.6 g·L^-1^ MS salts, 20 g·L^-1^ D-mannitol, 0.5 g·L^-1^ 2-(N-morpholino)-ethane sulfonic acid, 0.5 g·L^-1^ polyvinylpyrrolidone (40,000), 200 mg·L^-1^ L-glutamine, 40 mg·L^-1^ adenine, 0.1 mg·L^-1^ naphthalene acetic acid, 0.1 mg·L^-1^ 6-benzylaminopurine, 0.5 mg·L^-1^ nicotinic acid, 0.5 mg·L^-1^ pyridoxine, 2 mg·L^-1^ glycine, 2% sucrose, adjusted to pH 5.8, and 2 g·L^-1^ gelrite). The sterilized medium was supplemented with 1 mg·L^-1^ zeatin riboside, 0.1 mg·L^-1^ naphthalene acetic acid, and 200 mg·L^-1^ kanamycin. Untransformed potato leaves from the variety Desiree were included as controls. After one month, the presence or not of a callus confirmed or not the transgenic nature of the leaf.

### Testing for Bacterial wilt resistance

Transgenic events were tested against the *R. solanacearum* isolate CIP-204 (Bv2A Race 3, phylotype IIB sequevar 1) under greenhouse conditions in separate batches (referred to as experiments hereafter), with three plants per event. Events showing the least amount of disease and infection were selected for more extensive testing. Selected transgenic events were evaluated with 15 repetitions per event, placed in three separate trays (five plants per tray) and tested either with CIP-204 or CIP-475 (Bv2A Race 3, phylotype IIB sequevar 1) of *R. solanacearum*.

For each experiment, *in vitro* plantlets were transplanted into Jiffy-Strips 7 (Jiffy Products [NB] Ltd, Canada) containing 20 g of Promix Bx substrate: sand mixture) and maintained under greenhouse conditions at 18-20°C. Approximately two weeks later, when the plants were 10-15 cm long and had adequate roots, they were transplanted into 3.5-inch-diameter pots containing 85 g of promix Bx substrate (Premier Tech, Canada). 50 mL solution of fertilizer (N–P–K: 6-7-22) was applied at the time of planting. Plants were irrigated daily, except for one day before inoculation.

For plant inoculation, bacterial suspensions were prepared by culturing isolates for 48 h at 30°C in modified Kelman’s medium (MKM) lacking 2,3,5-triphenyl-tetrazolium chloride (TZC). Bacterial cells were harvested in sterile distilled water, and the bacterial concentration was determined by measuring the absorbance at 600 nm (OD_600_) using a spectrophotometer. The absorbance reading at OD_600_ = 0.1 corresponded to a bacterial concentration of 2 × 10^8^ cells·mL^-1^. Two weeks after transplanting, the plants were inoculated by pouring 40 mL of the bacterial suspension (final concentration of 4 × 10^7^ cells·g^-1^ soil) into the pots. Plants were maintained until one month after inoculation, during which temperatures ranged from 24 to 26°C. Non-transgenic potato varieties Desiree and Cruza 148 were used as susceptible and moderately resistant controls, respectively. The latter has been found to have a degree of resistance to bacterial wilt (French & Lindo, 1982).

For each transgenic event, wilt incidence was recorded twice a week, beginning when the first wilting symptom appeared in any plant, and continuing until wilting was complete. Stems of events that exhibited no wilt symptoms were analyzed to detect latent infections. An approximately 5-cm-long stem segment near the base of the plant was cut from symptomless plants, disinfected with 70% alcohol, and rinsed with sterile distilled water. The stem fragments were placed in plastic bags, weighed, and crushed using a wooden rolling pin. The samples were homogenized using 3 ml of sterile citrate extraction buffer (0.1 M citric acid, 0.1 M sodium citrate, pH 5.6) per gram of stem tissue. Stem extract was used to assess for the presence of *R. solanacearum* in post-enrichment NCM-ELISA tests (Priou *et al*., 1999), and positive samples were confirmed by isolating *R. solanacearum* on MKM medium with TZC (French *et al*., 1995).

Data were subjected to analysis of variance (ANOVA) using the F-test (p < 0.05). Differences in the level of resistance between the best events were measured as percentages of wilt, the area under the disease progress curve (AUDPC), the percentage of symptomless plants with latently infected stems, and the total percentage of infected plants (wilted plants + symptomless plants with latently infected stems) and analyzed using the Scott–Knott Clustering Algorithm for Single Experiments method detailed by Scott & Knott (1974), and the Kruskal-Wallis test and multiple comparisons of treatments (Conover, 1999), using the Agricola package (De Mendiburu, 2012) of R software (R Core Team, 2019). The AUDPC of the transgenic events was used to summarize the progression of disease severity over time and was compared to that of the untransformed controls: moderately resistant (Cruza 148) and susceptible (Desiree).

### Inoculation, sampling and processing for RT-qPCR

Gene expression analysis was performed on the leaves of the transgenic plants one month after inoculation with isolate CIP-204 inoculation with sterile water. For inoculation, the plant roots were damaged with a scalpel, and 40 mL of the bacterial suspension was immediately poured into the soil (final concentration of each isolate was 5 × 10^7^ cells·g^-1^ soil). The plants were maintained under the same conditions as those described above. Three leaf or root samples were collected as biological replicates, washed with distilled water, immediately frozen in liquid nitrogen, and stored at −70°C until processing.

These samples were collected on the 22nd day after inoculation to ensure sufficient root growth. Frozen leaf or root samples were ground with a mortar and pestle in liquid nitrogen, and 100 mg of the powder was used for total RNA extraction using TRIzol Reagent (Invitrogen^®^) following the manufacturer’s protocol. RNA extraction was performed in triplicate for each sample.

### Real-time quantitative PCR

The expression of the β*hpmeh* gene was assessed using real-time quantitative PCR (RT-qPCR) and the *ef1*α gene as reference gene (Nicot *et al*., 2005). RNA extraction was cleaned with DNase treatment using Turbo DNA-free (Ambion^®^), following the manufacturer’s protocol. The reaction mixture was prepared in a total volume of 10 µL containing 4 µL of template (50 ng), 0.4 µM of each primer, and Power SYBR^®^ Green PCR Master Mix (Applied 274 Biosystems^®^ StepOne Real-Time PCR System). The expression of the β*hpmeh* gene was determined using the primers (pameh150Fw 5’-GCT ATG TTC GCT GCC CAA GG-3’ and pameh150Rv 5’-CGT TCC TTC CGG TAC CAA CG-3’) and the *ef1*α gene was amplified using the primers ef1αFw 5’-ATT GGA AAC GGA TAT GCT CCA-3’ and ef1αRv 5’-TCC TTA CCT GAA CGC CTG TCA-3’ (Nicot *et al*., 2005). The following PCR program was used: 95°C for 10 min, 95°C for 15 sec, and 55°C for 1 min for 40 cycles, followed by 95°C for 15 sec, 60°C for 1 min, and 95°C for 15 sec (melting curve analysis). The *ef1*α gene was used to normalize the expression levels, and differential expression was determined using REST^®^ 2009 software. No Ct values were obtained for non-transgenic events, so Ct values were set to 40 cycles (the number of cycles run and thus the maximum theoretical detection limit) to enable comparison by relative expression levels. Differences in RT-qPCR results were tested for statistical significance by ANOVA and Tuckey’s multiple comparison test. The significance values (pH1) were calculated using the REST software, which evaluates the probability of the hypothesis that there are differences in gene expression between the analyzed groups (H1), is true.

## Results

### Bacterial wilt resistance evaluation

Three hundred and sixty transgenic events (197 events with *GRP1.8-pameh*, and 163 transgenic events with the *35S-pameh*) were produced and evaluated from explants infected with *Agrobacterium* containing the corresponding gene construct. All transgenic events were screened for BW resistance using isolate CIP-204 in five experiments (*Exp 1 to 5*) with 3 replicates per genotype (Figure 2). Of the 360 events tested, 24 did not develop any wilt symptoms, 174 had low to high wilting symptoms, whereas 162 were fully susceptible (Table S1). This large-scale screening allowed the elimination of transgenic events that were as susceptible as the control Desiree. The bioassay data were not further analyzed due to the small number of repetitions.

**Figure 2.**
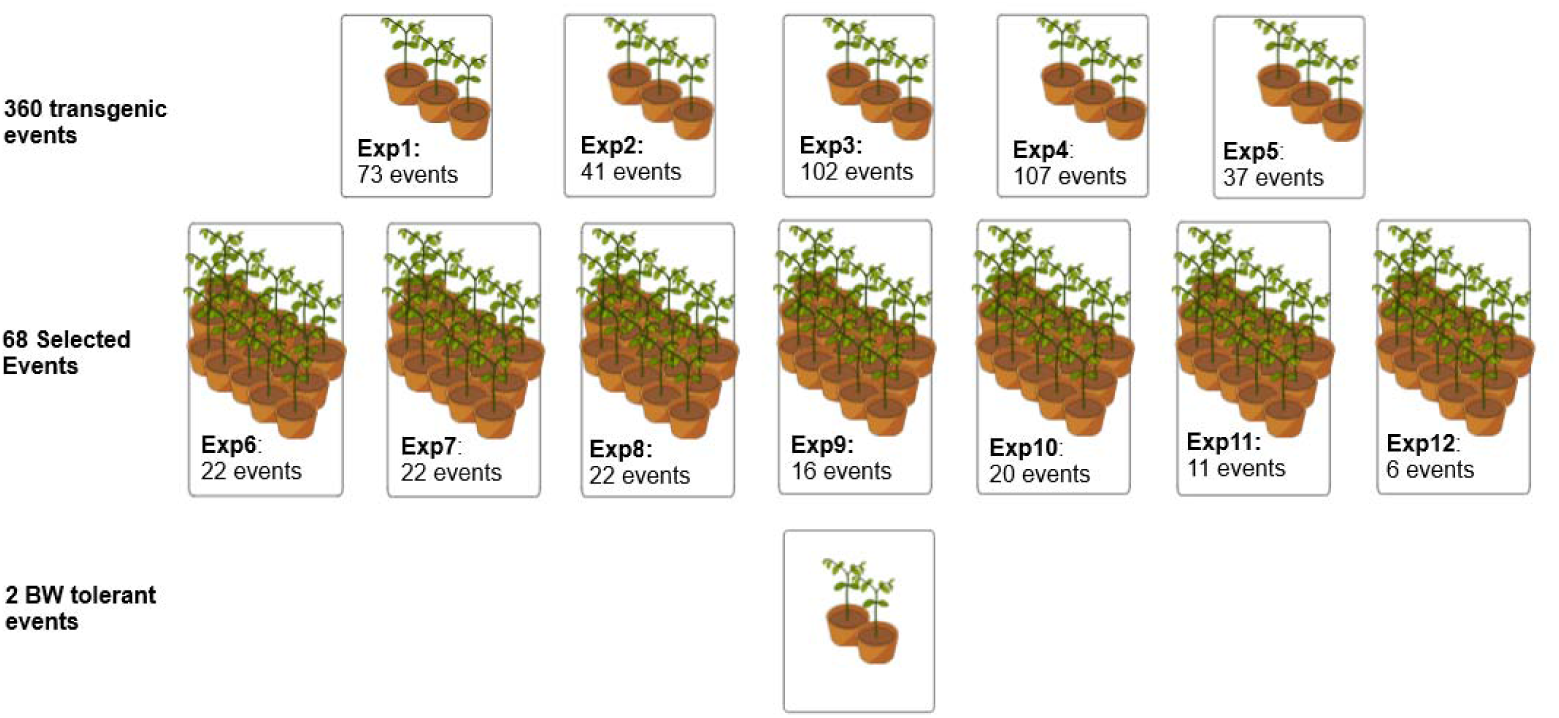
Schematic representation of screening transgenic events for BW resistance in a total of 12 experiments from an initial large-scale screening of 360 events in five experiments with (3 plants per genotype), followed by a screening of 68 selected transgenic events in seven experiments *Exp* 6 to 12 (15 plants per event) which resulted in identifying 22 highly confident resistant transgenic events of which 2 are as resistant as Cruza 148 to bacterial wilt disease, inoculated with CIP-204 or CIP-475 (*Exp* 8).

Sixty-eight transgenic events were selected from this large-scale screening. Sixty-two were selected based on a reduced incidence of wilt and no latent infection compared to the control. Four events with latent infections and two highly susceptible were included as well. The 68 events were evaluated in seven experiments (*Exp 6 to 12*) with 15 repetitions per genotype using isolate CIP 204 (Figure 2). Twenty-two of these transgenic events were additionally evaluated with another strain of *R. solanacearum*, strain CIP-475. No differences in virulence or aggressivity were observed between the two strains on the susceptible control variety Desiree. Bioassay data were thus combined.

In the experiments (*Exp 6-10*, *12*) with 15 plants, statistical analysis was performed to identify the most resistant plants; the detailed results are presented in the supplementary table (Table S2). The Scott–Knott test was used in *Exp 6*, *7*, *9*, and *10*. In *Exp 8*,*11* and *12*, the Kruskal-Wallis test was used. The latter test, a rank-based nonparametric test, was chosen because it can detect differences between groups with a small number of samples. The mean ranks values in Kruskal-Wallis are compared to determine if the differences are statistically significant among them. In all experiments, except for *Exp 8*, the resistant control Cruza 148 showed significantly reduced wilt, AUDPC, and total infected plants compared to the non-transgenic Desiree.

*Exp 6* (22 events) resulted in one resistant event (GRP 3.11) with 6.7% wilting symptoms, 6.7% latent infection (13.3% total infected plants) and the lowest AUDPC (113.3) values, similar to the moderately resistant control Cruza 148 (wilting: 6.7%, latent infection: 20%, total infected plants: 26.7%, and AUDPC: 100), followed by event II 4.16, with 53.3% of total infected plants. The bioassay data for these three genotypes were statistically significant.

The *Exp 7* (22 events) did not show any significant differences between events or controls in wilt and AUDPC results, but there was a significant reduction in the total percentage of infected plants (wilted + latently infected) for nine events compared to the non-transgenic Desiree. Because wilt incidence was relatively low in this experiment compared to the others, the conditions were likely unfavorable for disease development.

The *Exp 8* (22 events), with RSSC strain CIP-475, showed one event (GRP 3.11) with significantly reduced mean ranks of wilt incidence, AUDPC, and total infected plants compared to non-transgenic Desiree. Similar to the result from *Exp 6*, the event II 4.16 presented AUDPC values significantly lower than those of Desiree. When analyzing the total % of infected plants, the events GRP 3.11, GRP 2.18, GRP 2.3, GRP 2.6, and GRP 1.5, had significantly lower infection rates than Desiree.

The *Exp 9* (16 events) showed five events (GRP 7.28, GRP 7.15, GRP 9.7, GRP 6.48, and GRP 5.14) with significantly reduced total infected plants compared to Desiree.

The *Exp 10* (20 events) resulted in four events (II 10.29, II 4.23, II 8.5, and II 5.8) with significantly reduced wilt percentage and AUDPC (except for II 5.8), two of which (II 5.8, and II 10.29) were also reduced in total infected plants.

The *Exp 11* (10 events) resulted in one event (II 10.31) with significantly reduced mean ranks of wilt, AUDPC or infection, including the control Cruza 148. However, the Desiree control did not behave as expected and was not among the most wilted genotypes. Hence, the analysis using relative AUDPC to Desiree did not include the data from *Exp 11*.

The *Exp 12* (six events) resulted in two events (II 6.3 and II 4.16) with significantly reduced mean ranks of AUDPC and wilt percentage, and one event II 10.29 as resistant in terms of total wilt percentage, compared to Desiree.

Twenty-two transgenic events were finally selected from the second level screening based on the criteria of having gone through at least three independent BW resistance evaluations (experiments). Indeed, wilting symptoms varied significantly from experiment to experiment as evidenced from the bioassay data of the controls. One of the experiments (*Exp 11*) has not been included in the BW resistance analysis because the Desiree control did not perform as expected. We therefore selected further our sample of transgenic events for those with at least three bioassay datasets that supported significant increase in resistance, which resulted in 22 transgenic events with highly confident bioassay data (Table 1).

**Table 1.**
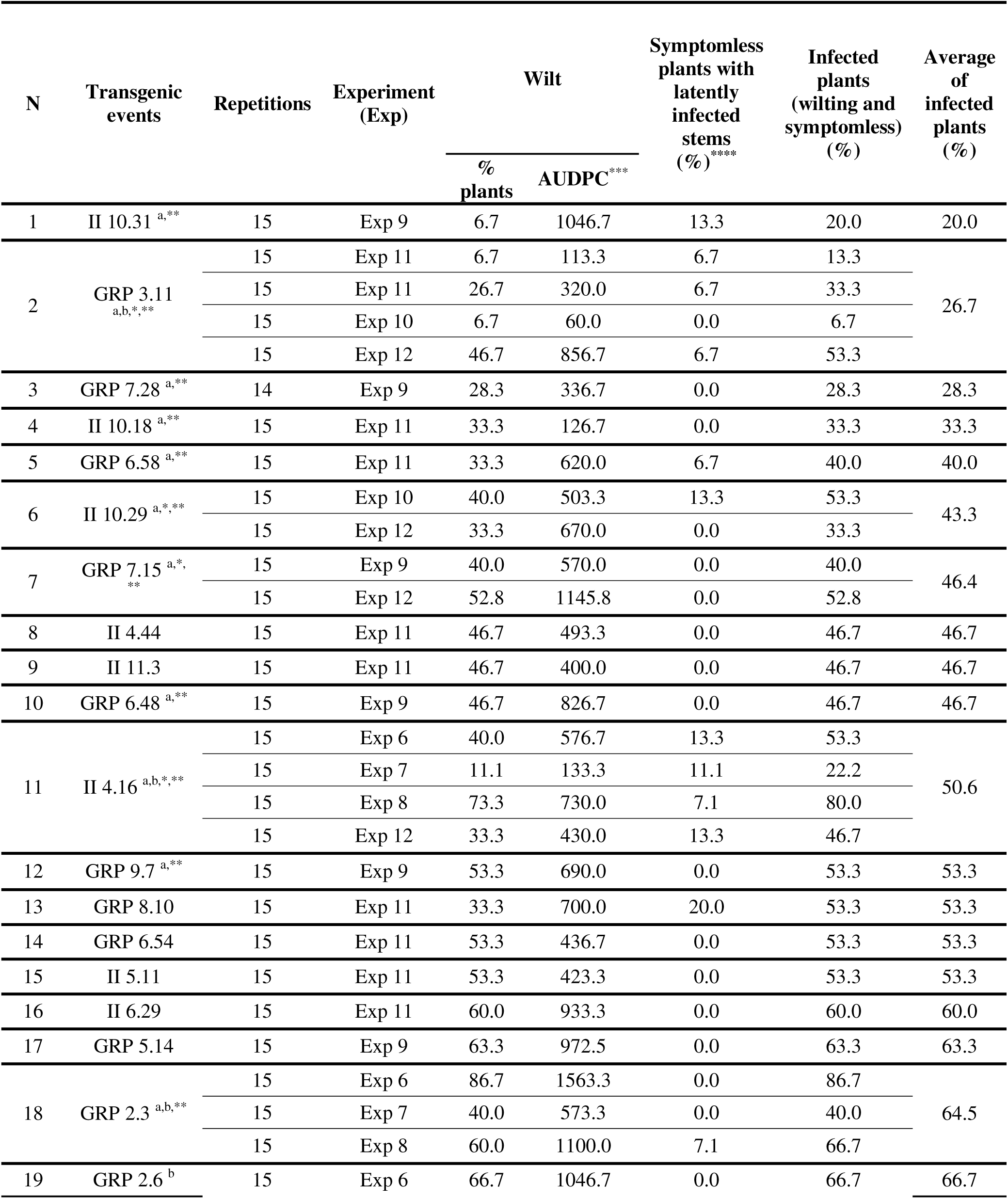
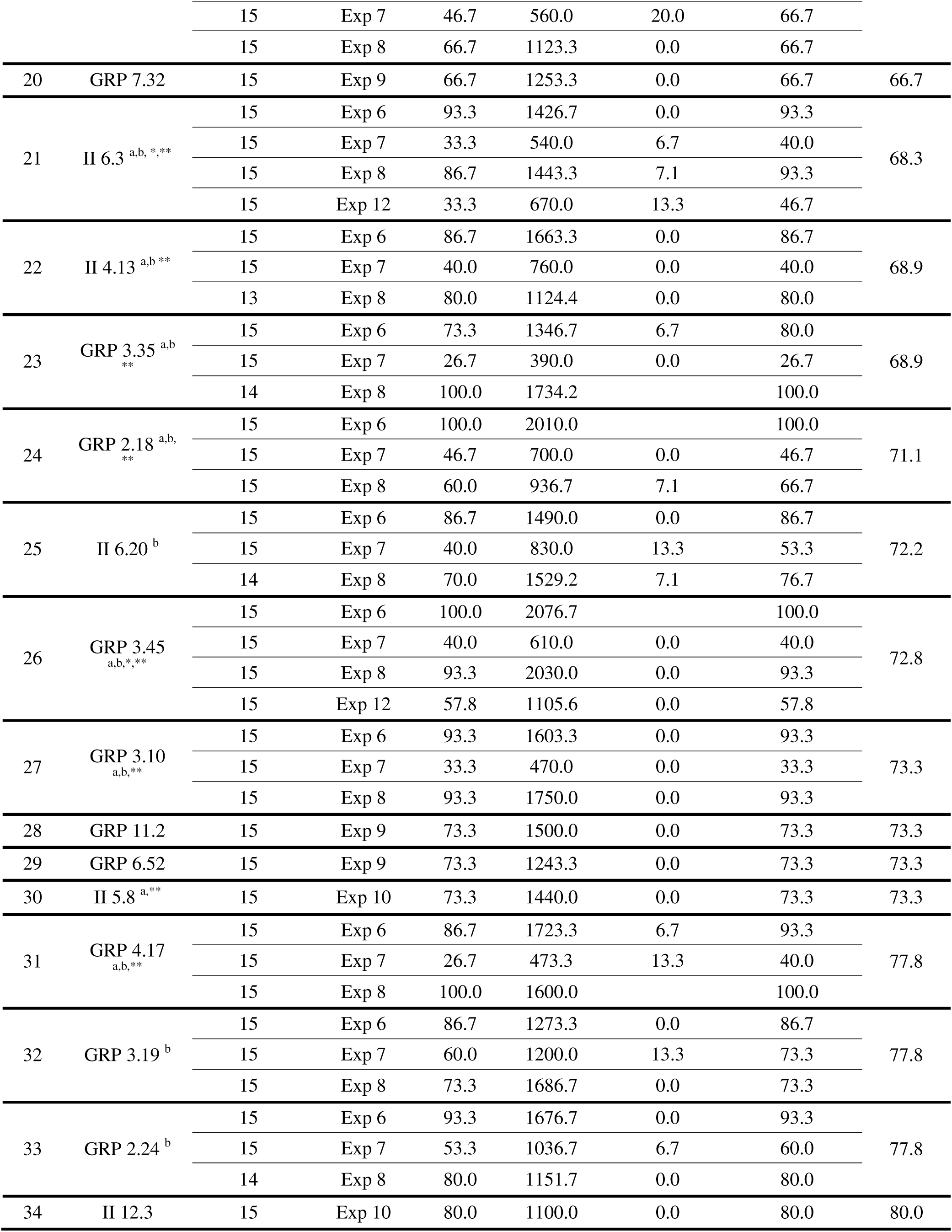
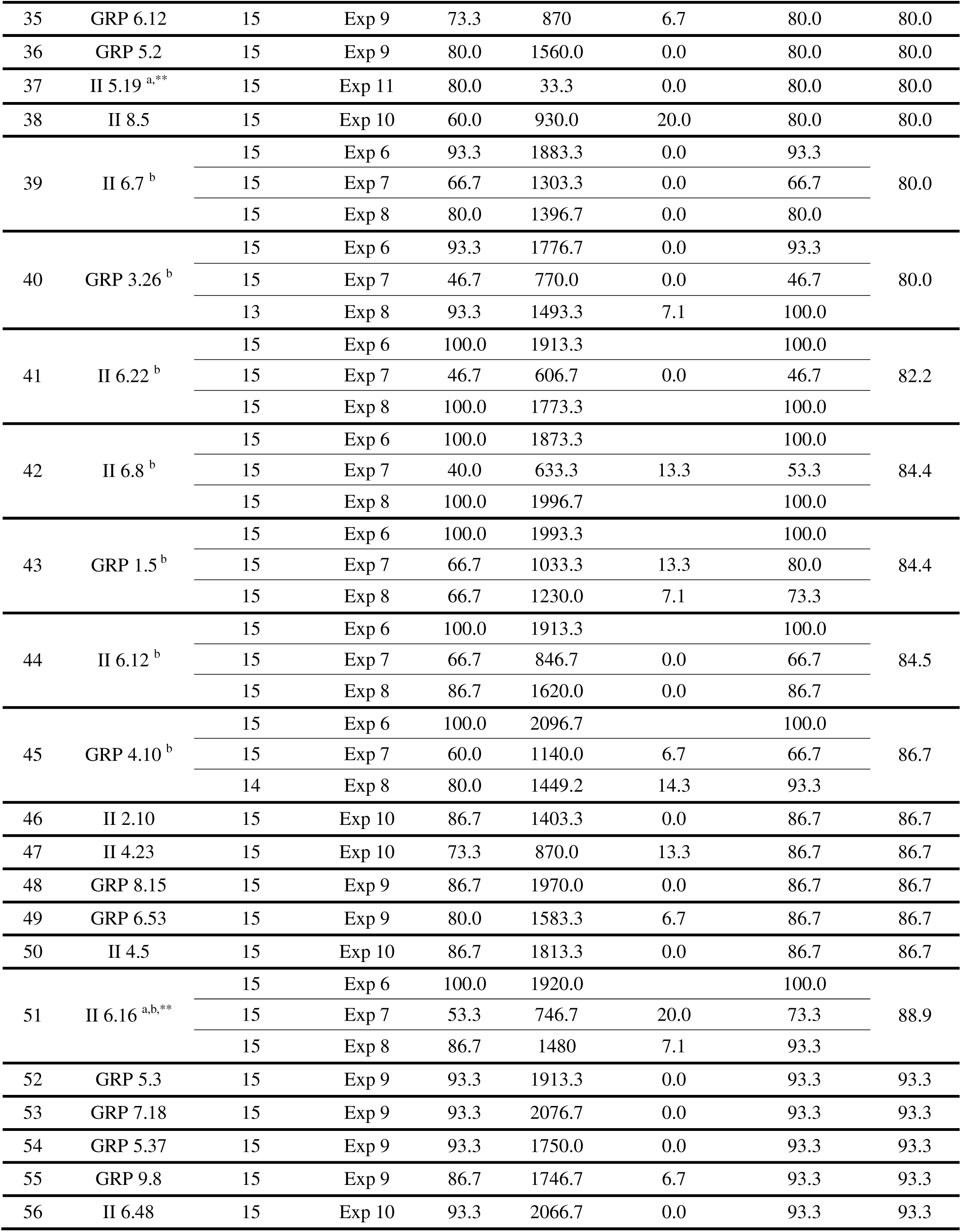
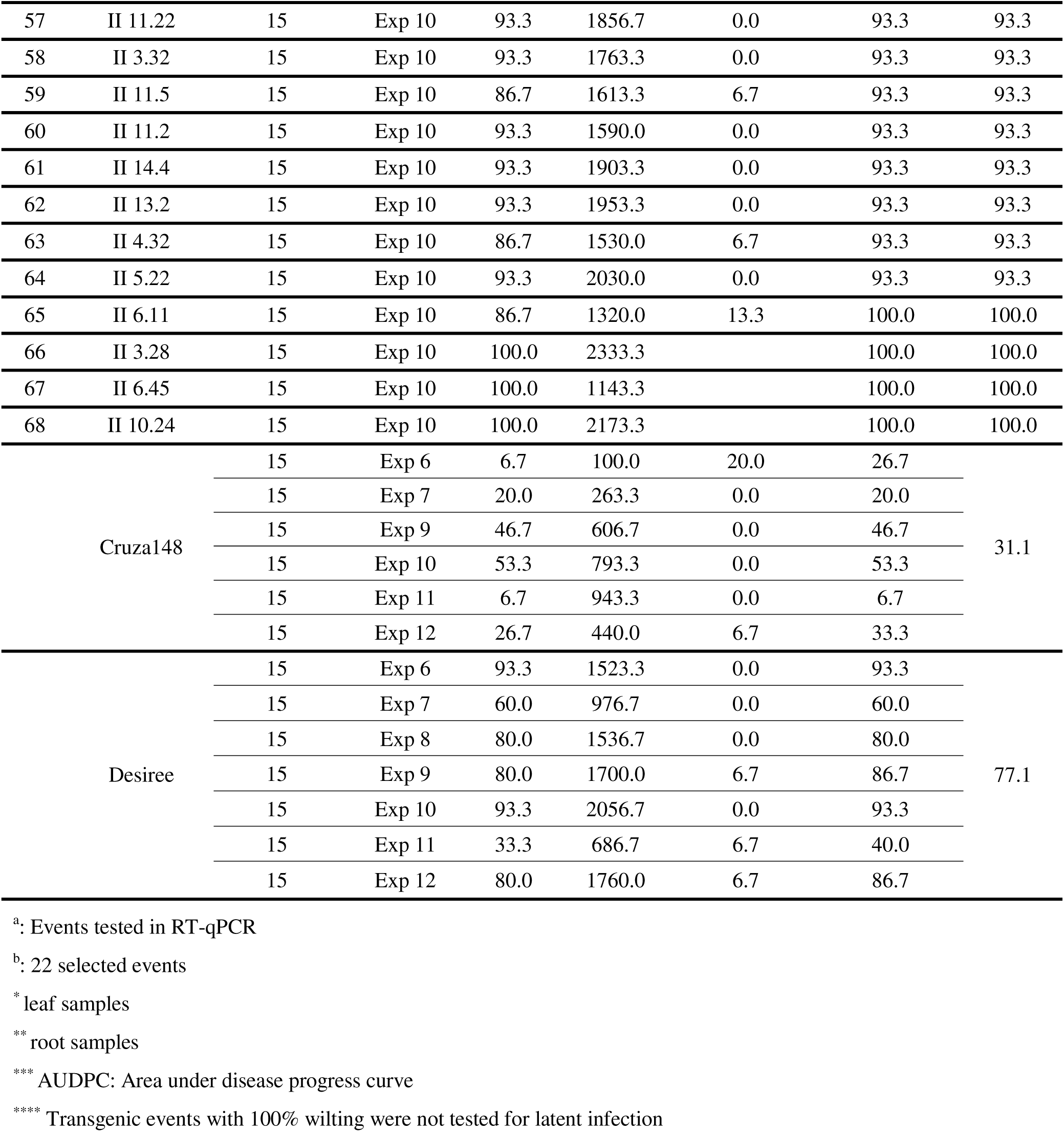
List of transgenic events (n=68), and 2 controls that were tested more than once (sorted by the average of total infected plants).

The further analysis of BW resistance assay data conducted on the 22 transgenic events with highly confident BW resistance data resulted in the identification of 2 transgenic events, GRP 3.11 and II 4.16 with an average AUDPC relative to the AUDPC of the susceptible control Desiree of 23% and 31% respectively (Table 1). This is similar to the value of the moderately BW resistant variety Cruza 148 (27%). Most of the rest of the transgenic events were almost as susceptible as Desiree (Figure 3). A statistical analysis was conducted on the bioassay data and is presented hereafter.

**Figure 3.**
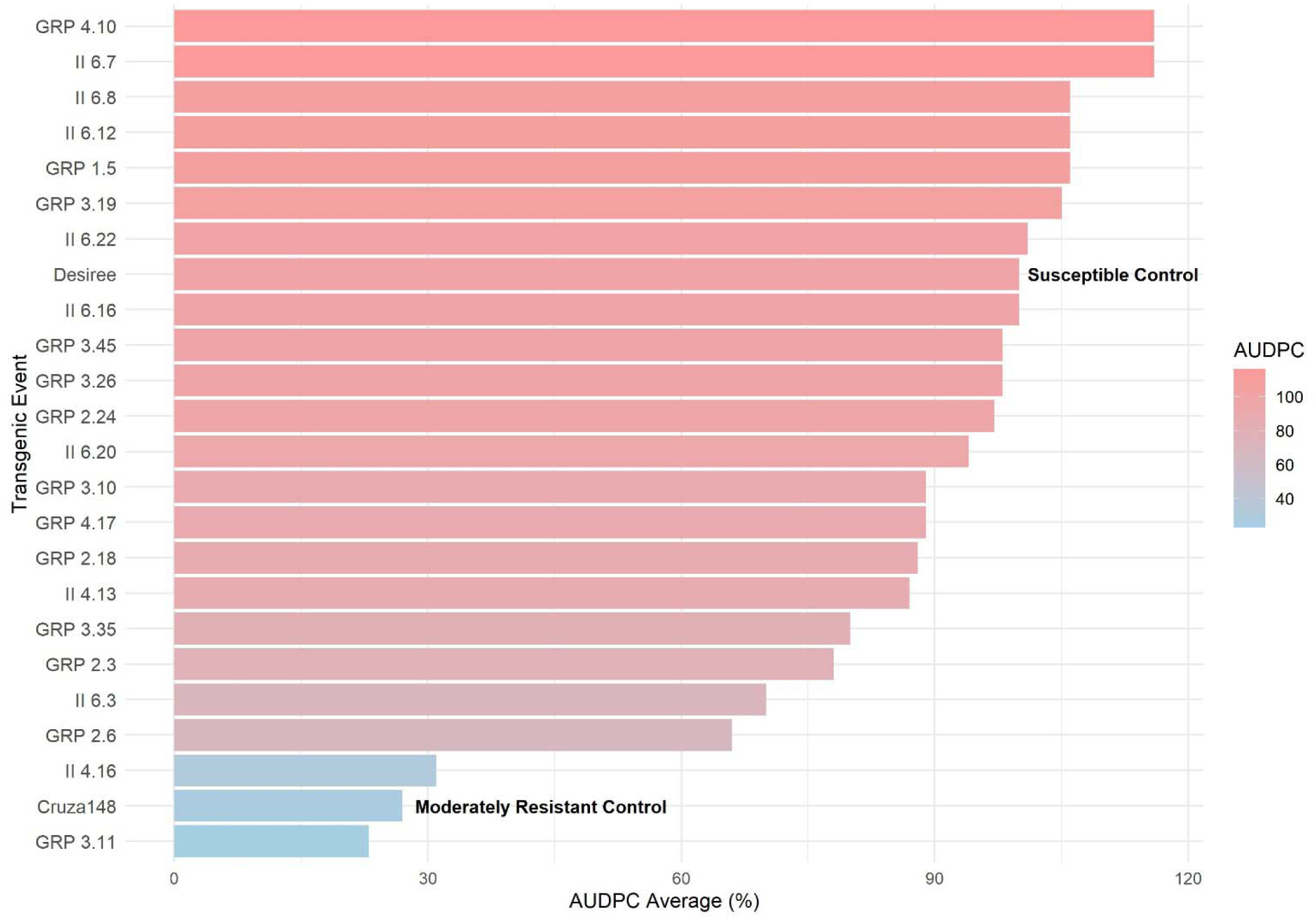
Disease progression across transgenic events based on the average area under disease progress curve (AUDPC) from *Exp 6-10, Exp 12*. Desiree and Cruza 148 were used as susceptible and moderately resistant controls, respectively.

### βhpmeh gene expression

RNA was extracted from leaves from plants one month after inoculation from *Exp* 12 (GRP 3.45, GRP 3.11, II 4.16, II 6.3, GRP 7.15, and II 10.29) and were analyzed using RT-qPCR. Expression levels from transgenic plants were analyzed relative to GRP 7.15, chosen because of its lowest expression level in the experiment (Figure 4). All measurements were normalized to *ef1*α before performing relative expression comparisons. Events II 6.3, II 4.16, and II 10.29 were identified as the events with highest levels of β*hpmeh* expression and correlated with lowest level of wilting in the same plants, and II.6.3 had a significantly higher level of expression than the rest. Also, the pH1 values (<0.05) support that this event (as well as II.10.29 and II.4.16) had significantly higher expression compared to the non-transgenic control (Figure 4).

**Figure 4.**
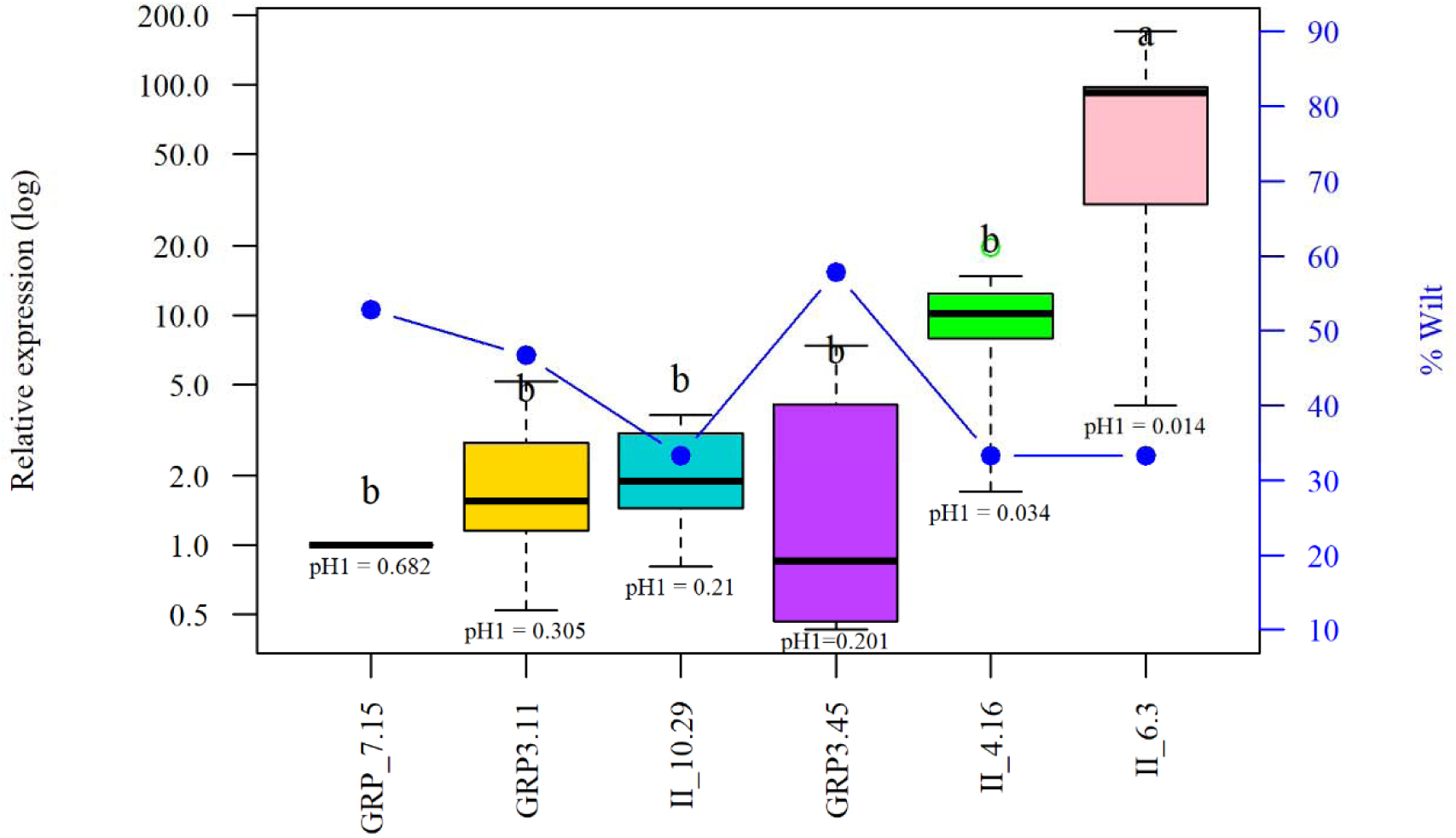
Box plot of the relative expression of β*hpmeh* gene in leaves of transgenic events relative to GRP 7.15 (value=1) one month after inoculation with CIP-204 (experiment 12). Expression levels were normalized to those of *ef1*α. The phenotypic results (%) are plotted on the 2nd axis. Statistically significant differences in expression between different events, as determined by Tukey’s multiple comparison test (α: 0.05), are shown by different letters above each box plot. The significance values (pH1) for expression as compared to non-transgenic Desiree are shown below each box plot.

Additionally, we analyzed the relative expression of β*hpmeh* in root samples from transgenic events 22 days after inoculation with CIP204, using GRP 2.18 as a reference (Figure 5). This experiment was conducted independently (Experiment 13) due to the use of a distinct tissue for expression analysis. A subset of 9 randomly selected transgenic events (II 10.31, GRP 9.7, II 5.19, GRP 6.48, II 5.8, GRP 7.28, GRP 6.58, II 4.13, and II 10.18) that were not part of the twenty-two transgenic events previously selected was included. Overall, the analysis comprised 12 events harboring the GRP promoter (GRP 3.45, GRP 3.10, GRP 2.3, GRP 4.17, GRP 2.18, GRP 3.35, GRP 7.15, GRP 6.48, GRP 6.58, GRP 9.7, GRP 3.11, and GRP 7.28) and 9 events harboring the 35S promoter (II 5.8, II 6.16, II 4.13, II 6.3, II 10.29, II 5.19, II 4.16, II 10.31, and II 10.18).

**Figure 5.**
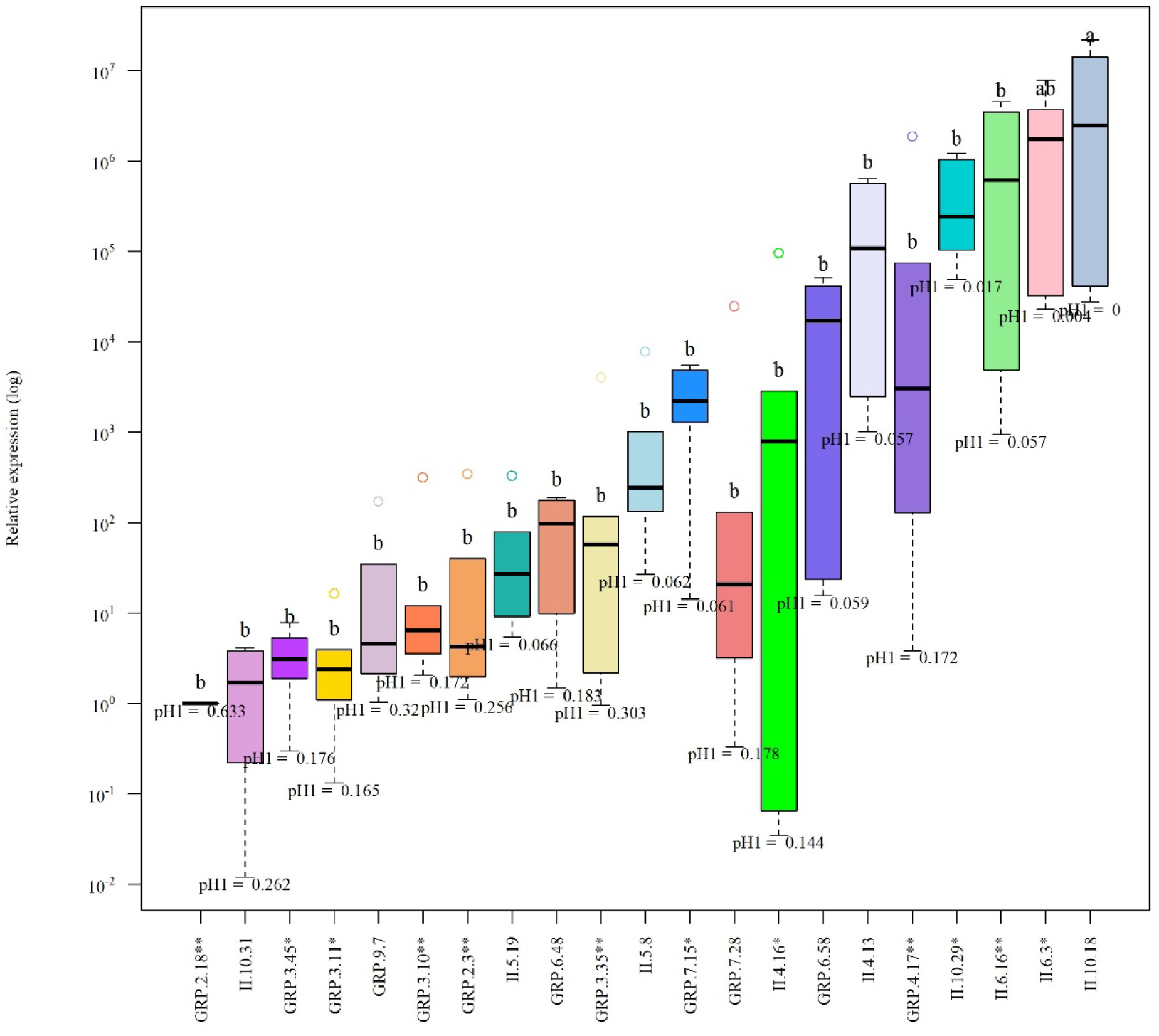
Relative expression of the β*hpmeh* gene in roots of transgenic events 22 days after inoculation with CIP204, relative to GRP 2.18 (colored whisker plots). The expression levels were normalized to those of *ef1-a*. Non-transgenic ‘Desiree’ was used as a susceptible control. Statistically significant differences in expression between different events, as determined by Tukey’s multiple comparison test (α: 0.05), are shown by different letters above each box plot. *Events included in *Exp* 12 and in the 22 selected events, **Events belonging to the 22 selected events. The significance values (pH1) for expression as compared to non-transgenic ‘Desiree’ are shown below each box plot.

Since root sampling was destructive, phenotypic assessment could not be performed on the same plants. Among the tested events, II 10.18 (excluded from Experiment 12 due to logistical reasons) exhibited the highest relative expression.

## Discussion

In this study, we evaluated the ability of the β*hpmeh* gene, introduced into potato, to confer resistance to bacterial wilt following artificial inoculation with *R. solanacearum*. Two gene constructs were tested: one driven by the root-specific GRP 1.8 promoter from bean and the other by the constitutive CaMV 35S promoter. The highly susceptible cultivar Desiree was used to generate 360 transgenic events, and resistance was assessed in 12 independent bioassays, each including the moderately resistant cultivar Cruza 148 as a reference.

Screening for bacterial wilt resistance is challenging due to variability in pathogen aggressiveness and difficulties in standardizing environmental conditions such as temperature, humidity, and soil-related factors. For this reason, bioassay data were analyzed separately for each experiment rather than pooled. After large-scale screening of 360 transgenic events, 68 were selected for further evaluation of BW resistance. Of these, 22 events evaluated in at least three experiments were considered to have reliable resistance data. Two events, driven by either promoter, exhibited resistance levels comparable to the moderately resistant control Cruza 148. A few events showed intermediate resistance between Cruza 148 and the susceptible cultivar Desiree, whereas the majority remained as susceptible as Desiree. In one experiment, β*hpmeh* gene expression in leaves was positively correlated with resistance levels. However, root expression levels did not differ significantly between the GRP1.8 and CAMV 35S promoter constructs.

These results are consistent with previous studies targeting quorum sensing in *R. solanacearum* to reduce disease. For instance, Wang *et al*. (2023) demonstrated that *Pseudomonas* strains capable of degrading 3-OH PAME effectively suppressed virulence factors of *R. solanacearum* in *C. equisetifolia*, peanut, and tomato plants. Achari and Ramesh (2015) reported that bacteria such as *Stenotrophomonas maltophilia* and *Rhodococcus corynebacterioides,* which possess 3-OH PAME degrading activity, reduced bacterial wilt incidence in eggplant seedlings, although the genes involved in these enzymatic activities remain unidentified.

Given that some transgenic events achieved resistance levels comparable to Cruza 148 under controlled conditions, it is reasonable to expect similar moderate resistance under field conditions. However, from an epidemiological perspective, moderate resistance may pose a risk, as plants can act as asymptomatic carriers of *R. solanacaearum*, facilitating disease spread. This phenomenon has been documented in Ethiopia, where latent infections contributed to significant spread and subsequent losses in potato production (Abdurahman *et al*., 2017). Therefore, combining the β*hpmeh* gene with additional BW resistance genes may represent a more effective strategy to achieve durable resistance while minimizing the risk of latent infection associated with vegetatively propagated seed.

One promising candidate for gene pyramiding is the *Arabidopsis thaliana <u>EF</u>*-Tu *r*eceptor (*EFR*), a receptor kinase that activates immune signaling pathways in response to bacterial elicitors (Zipfel *et al.,* 2006). Transgenic potato plants expressing the *EFR* gene have shown enhanced BW resistance under controlled conditions (Boschi *et al*., 2017). Targeting infection sites through quorum-quenching and/or pathogen-associated molecular pattern (PAMP) recognition may provide synergistic effects, potentially leading to enhanced resistance.

However, the number of identified *R. solanacearum* PAMPs and corresponding plant pattern recognition receptors (PRRs) remains limited. Expanding this knowledge will be essential for developing more effective resistance strategies (An & Zhang, 2024). Transgenic expression of several immune receptors from *Nicotiana benthamania* has shown promising results in enhancing resistance to bacterial pathogens (Schultink *et al*., 2017, 2019). Gene stacking approaches combining *EFR*, *Roq1*, and *Jim2* are currently under investigation for conferring complete resistance to bacterial wilt (Mwangi *et al*., 2024).

Understanding the interactions among quorum-sensing signals, PAMPs, and PRRs is essential for developing effective and durable resistance strategies against *R. solanacearum* in crop plants (Huet, 2014; An & Zhang, 2024).

In conclusion, this study provides evidence that quorum quenching mediated by the βHPMEH enzyme represents a promising strategy for controlling bacterial wilt in potato. Compared with conventional antibacterial approaches, quorum quenching may reduce the likelihood of resistance development, as it targets bacterial communication systems rather than essential cellular processes (Wang et al., 2022).

## Author contributions

1. M. Izarra: Data curation, Methodology, Formal analysis, Investigation, Writing – original draft, Writing – review & editing.
2. L. Gutarra: Supervision, Data curation, Methodology, Formal analysis, Investigation.
3. E. Fernandez: Data curation, Methodology, Formal analysis, Investigation
4. E. Huaman: Investigation
5. M. Ghislain: Formal analysis, Writing – review & editing.

JF. Kreuze: Conceptualization, Project administration, Supervision, Formal analysis, Investigation, Methodology, Writing – review & editing.

## Acknowledgements

We gratefully acknowledge technical support from Yuan Guo and José Rodriguez.

## Funding

This work was funded by the CGIAR consortium research program on roots, tubers and bananas and the CGIAR initiative on Plant Health, both supported by the donors of the CGIAR Trust Fund (http://www.cgiar.org/funders/).

## Conflict of Interest

The authors declare that they have no known competing financial interests or personal relationships that could have appeared to influence the work reported in this paper.

## Data availability statement

Not applicable.

## Ethics declaration

Not applicable.

## Notes

### Competing Interest Statement

The authors have declared no competing interest.

### Summary of Updates

The manuscript has been revised in accordance with the editorial comments and journal guidelines. The Introduction was substantially rewritten to improve originality and clarity, and additional edits were made throughout the manuscript to improve language, formatting, and consistency. The study conclusions remain unchanged.

## References

Abdurahman A, Griffin D, Elphinstone J et al., 2017. Molecular characterization of Ralstonia solanacearum strains from Ethiopia and tracing potential source of bacterial wilt disease outbreak in seed potatoes. Plant Pathology 66, 826–834.

Achari GA, Ramesh R, 2015. Characterization of bacteria degrading 3-hydroxy palmitic acid methyl ester (3OH-PAME), a quorum sensing molecule of Ralstonia solanacearum. Letters in Applied Microbiology 60.

Amack SC, Antunes MS, 2020. CaMV35S promoter – A plant biology and biotechnology workhorse in the era of synthetic biology. Current Plant Biology 24, 100179.

An Y, Zhang M, 2024. Advances in understanding the plant-*Ralstonia solanacearum* interactions: Unraveling the dynamics, mechanisms, and implications for crop disease resistance. New Crops 1, 100014.

Boschi F, Schvartzman C, Murchio S et al., 2017. Enhanced Bacterial Wilt Resistance in Potato Through Expression of Arabidopsis EFR and Introgression of Quantitative Resistance from Solanum commersonii. Frontiers in Plant Science 8.

Brumbley SM, Carney BF, Denny TP, 1993. Phenotype conversion in Pseudomonas solanacearum due to spontaneous inactivation of PhcA, a putative LysR transcriptional regulator. Journal of Bacteriology 175, 5477–5487.

CABI, 2023. *Ralstonia solanacearum* (bacterial wilt of potato). Cabi Digital Library.

Champoiseau PG, Jones JB, Allen C, 2009. Ralstonia solanacearum Race 3 Biovar 2 Causes Tropical Losses and Temperate Anxieties. Plant Health Progress 10, 35.

Charkowski A, Sharma K, Parker ML, Secor GA, Elphinstone J, 2020. Bacterial Diseases of Potato. In: Campos H, Ortiz O, eds. The Potato Crop: Its Agricultural, Nutritional and Social Contribution to Humankind. Cham: Springer International Publishing, 351–388.

Chesneau T, Maignien G, Boyer C et al., 2018. Sequevar Diversity and Virulence of Ralstonia solanacearum Phylotype I on Mayotte Island (Indian Ocean). Frontiers in Plant Science 8.

Clough SJ, Lee KE, Schell MA, Denny TP, 1997a. A two-component system in Ralstonia (Pseudomonas) solanacearum modulates production of PhcA-regulated virulence factors in response to 3-hydroxypalmitic acid methyl ester. Journal of Bacteriology 179, 3639–3648.

Conover WJ, 1999. Practical nonparametric statistics. New York: Wiley.

De Mendiburu F, 2012. Agricolae:[: Statistical Procedures for Agricultural Research.

Denny TP, Hayward AC, 2001. Gram negative bacteria: Ralstonia. In: Schaad NW, Jones JB, Chun W, eds. Laboratory Guide for Identification of Plant Pathogenic Bacteria. St. Paul, MN: APS Press, 51–74.

Dong Y-H, Wang L-H, Xu J-L, Zhang H-B, Zhang X-F, Zhang L-H, 2001. Quenching quorum-sensing-dependent bacterial infection by an N-acyl homoserine lactonase. Nature 411, 813–817.

Dong Y-H, Xu J-L, Li X-Z, Zhang L-H, 2000. AiiA, an enzyme that inactivates the acylhomoserine lactone quorum-sensing signal and attenuates the virulence of Erwinia carotovora. Proceedings of the National Academy of Sciences 97, 3526–3531.

Fegan M, Prior P, 2005. How Complex is the *Ralstonia Solanacearum* Species Complex. In: Allen C, Prior P, Hayward AC, eds. Bact Wilt Dis Ralstonia Solanacearum Species Complex. St. Paul, MN: American Phytopathological Society, 449–462.

Fernández E, Gutarra L, Kreuze J, 2015. Evaluación del gen que codifica la enzima βHPMEH para la inhibición de la marchitez bacteriana causada por Ralstonia solanacearum. Revista peruana de biología 22, 193–198.

Ferreira V, Pianzzola MJ, Vilaró FL et al., 2017. Interspecific Potato Breeding Lines Display Differential Colonization Patterns and Induced Defense Responses after *Ralstonia solanacearum* Infection. Frontiers in Plant Science 8.

Flavier AB, Clough SJ, Schell MA, Denny TP, 1997a. Identification of 3-hydroxypalmitic acid methyl ester as a novel autoregulator controlling virulence in *Ralstonia solanacearum*. Molecular Microbiology 26, 251–259.

Flavier AB, Ganova-Raeva LM, Schell MA, Denny TP, 1997b. Hierarchical autoinduction in Ralstonia solanacearum: control of acyl-homoserine lactone production by a novel autoregulatory system responsive to 3-hydroxypalmitic acid methyl ester. Journal of Bacteriology 179, 7089–7097.

French E, Gutarra L, Aley P, 1995. Culture media for *Ralstonia solanacearum* Isolation, identification and maintenance. Fitopatologia 30, 126–130.

French E, Lindo D, 1982. Resistance to Pseudomonas solanacearum in potato: specificity and temperature sensitivity.

Genin S, 2010. Molecular traits controlling host range and adaptation to plants in Ralstonia solanacearum. New Phytologist 187, 920–928.

Ghorai AK, Dutta S, Barman AR, 2022. Genetic diversity of Ralstonia solanacearum causing vascular bacterial wilt under different agro-climatic regions of West Bengal, India. PLOS ONE 17, e0274780.

Hayward AC, 1994. Systematics and phylogeny of Pseudomonas solanacearum and related bacteria. In: Hayward AC, Hartman GL, eds. Bacterial Wilt: the Disease and its Causative Agent, Pseudomonas solanacearum. 123–135.

Hikichi Y, Mori Y, Ishikawa S et al., 2017. Regulation Involved in Colonization of Intercellular Spaces of Host Plants in Ralstonia solanacearum. Frontiers in Plant Science 8.

Hikichi Y, Yoshimochi T, Tsujimoto S et al., 2007. Global regulation of pathogenicity mechanism of *Ralstonia solanacearum*. Plant Biotechnology 24, 149–154.

Huet G, 2014. Breeding for resistances to *Ralstonia solanacearum*. Frontiers in Plant Science 5.

Keller B, Baumgartner C, 1991. Vascular-specific expression of the bean GRP 1.8 gene is negatively regulated. The Plant Cell 3, 1051–1061.

Lv X, Song X, Rao G et al., 2009. Construction vascular-specific expression bi-directional promoters in plants. Journal of Biotechnology 141, 104–108.

Medina-Bolivar F, Wright R, Funk V et al., 2003. A non-toxic lectin for antigen delivery of plant-based mucosal vaccines. Vaccine 21, 997–1005.

Mwangi MN, Witek AI, Magembe EM, Schultink A, Jones JD, Ghislain M, 2024. Development of Bacterial Wilt Resistant Potato Using Efr, Roq1 and Jim2 GENE. In: Plant and Animal Genome Conference/PAG31. PAG

Nicot N, Hausman J-F, Hoffmann L, Evers D, 2005. Housekeeping gene selection for real-time RT-PCR normalization in potato during biotic and abiotic stress. Journal of Experimental Botany 56, 2907–2914.

Paudel S, Dobhal S, Alvarez AM, Arif M, 2020. Taxonomy and Phylogenetic Research on Ralstonia solanacearum Species Complex: A Complex Pathogen with Extraordinary Economic Consequences. Pathogens 9, 886.

Peeters N, Carrère S, Anisimova M, Plener L, Cazalé A-C, Genin S, 2013. Repertoire, unified nomenclature and evolution of the Type III effector gene set in the Ralstonia solanacearum species complex. BMC Genomics 14, 859.

Priou S, Gutarra L, Aley P, 1999. Highly sensitive detection of *Ralstonia solanacearum* in latently infected potato tubers by post[enrichment enzyme[linked immunosorbent assay on nitrocellulose membrane. EPPO Bulletin 29, 117–125.

R Core Team, 2019. R: A language and environment for statistical computing.

Safni I, Cleenwerck I, De Vos P, Fegan M, Sly L, Kappler U, 2014. Polyphasic taxonomic revision of the Ralstonia solanacearum species complex: proposal to emend the descriptions of Ralstonia solanacearum and Ralstonia syzygii and reclassify current R. syzygii strains as Ralstonia syzygii subsp. syzygii subsp. nov., R. solanacearum phylotype IV strains as Ralstonia syzygii subsp. indonesiensis subsp. nov., banana blood disease bacterium strains as Ralstonia syzygii subsp. celebesensis subsp. nov. and R. solanacearum phylotype I and III strains as Ralstonia pseudosolanacearum sp. nov. International Journal of Systematic and Evolutionary Microbiology 64, 3087–3103.

Saile E, McGarvey JA, Schell MA, Denny TP, 1997. Role of Extracellular Polysaccharide and Endoglucanase in Root Invasion and Colonization of Tomato Plants by Ralstonia solanacearum. Phytopathology^TM^ 87, 1264–1271.

Schell MA, 2000. Control of Virulence and Pathogenicity Genes of Ralstonia Solanacearum by an Elaborate Sensory Network. Annual Review of Phytopathology 38, 263–292.

Schultink A, Qi T, Bally J, Staskawicz B, 2019. Using forward genetics in Nicotiana benthamiana to uncover the immune signaling pathway mediating recognition of the Xanthomonas perforans effector XopJ4. The New Phytologist 221, 1001–1009.

Schultink A, Qi T, Lee A, Steinbrenner AD, Staskawicz B, 2017. Roq1 mediates recognition of the Xanthomonas and Pseudomonas effector proteins XopQ and HopQ1. The Plant Journal: For Cell and Molecular Biology 92, 787–795.

Scott AJ, Knott M, 1974. A Cluster Analysis Method for Grouping Means in the Analysis of Variance. Biometrics 30, 507.

Shinohara M, Nakajima N, Uehara Y, 2007. Purification and characterization of a novel esterase (beta-hydroxypalmitate methyl ester hydrolase) and prevention of the expression of virulence by Ralstonia solanacearum. Journal of Applied Microbiology 103, 152–162.

Valls M, Genin S, Boucher C, 2006. Integrated Regulation of the Type III Secretion System and Other Virulence Determinants in Ralstonia solanacearum. PLOS Pathogens 2, e82.

Wang S, Hu M, Chen H et al., 2023. Pseudomonas forestsoilum sp. nov. and P. tohonis biocontrol bacterial wilt by quenching 3-hydroxypalmitic acid methyl ester. Frontiers in Plant Science 14, 1193297.

Wang H, Lin Q, Dong L et al., 2022. A Bacterial Isolate Capable of Quenching Both Diffusible Signal Factor- and N-Acylhomoserine Lactone-Family Quorum Sensing Signals Shows Much Enhanced Biocontrol Potencies. Journal of Agricultural and Food Chemistry 70, 7716–7726.

Whitehead NA, Barnard AM, Slater H, Simpson NJ, Salmond GP, 2001. Quorum-sensing in Gram-negative bacteria. FEMS microbiology reviews 25, 365–404.

Wu D, Ding W, Zhang Y, Liu X, Yang L, 2015. Oleanolic Acid Induces the Type III Secretion System of Ralstonia solanacearum. Frontiers in Microbiology 6.

Zhang L-H, 2003. Quorum quenching and proactive host defense. Trends in Plant Science 8, 238–244.

Zipfel C, Kunze G, Chinchilla D et al., 2006. Perception of the Bacterial PAMP EF-Tu by the Receptor EFR Restricts Agrobacterium-Mediated Transformation. Cell 125, 749–760.

