## Supplementary material for "Expression of the β*hpmeh* gene in transgenic events of the potato variety Desiree increases resistance to bacterial wilt caused by *Ralstonia solanacearum*": TableS1

**Table S1**. Percentage of wilt, percentage of symptomless plants with latently infected stems, and percentage of total infected plants (wilted plants and symptomless plants with latently infected stems) in transgenic events after soil drench inoculation with *Ralstonia solanacearum* at a concentration of 4 x $10^7$cells·$g^{-1}$ soil. The first, second, third, fourth, and fifth Experiments (three plants per event) [360 transgenic events and nine controls].

| N | Event | Promoter | Experiment (Exp) | plants | Wilting (%) | AUDCP* | Infection latent (%) ** | Infected plants (%) |
|---|---|---|---|---|---|---|---|---|
| | Desiree | Control | Exp 1 | 3 | 100.0 | 1566.7 | | 100.0 |
| 1 | GRP 1.1 | GRP1.8 | Exp 1 | 3 | 66.7 | 833.3 | 0.0 | 66.7 |
| 2 | GRP 1.7 | GRP1.8 | Exp 1 | 3 | 100.0 | 1283.3 | | 100.0 |
| 3 | GRP 2 | GRP1.8 | Exp 1 | 3 | 100.0 | 816.7 | | 100.0 |
| 4 | GRP 2.10 | GRP1.8 | Exp 1 | 3 | 33.3 | 333.3 | 33.3 | 66.7 |
| 5 | GRP 2.13 | GRP1.8 | Exp 1 | 3 | 66.7 | 1066.7 | 0.0 | 66.7 |
| 6 | GRP 2.18 | GRP1.8 | Exp 1 | 3 | 0.0 | 0.0 | 0.0 | 0.0 |
| 7 | GRP 2.19 | GRP1.8 | Exp 1 | 3 | 66.7 | 866.7 | 0.0 | 66.7 |
| 8 | GRP 2.2 | GRP1.8 | Exp 1 | 3 | 100.0 | 1100.0 | | 100.0 |
| 9 | GRP 2.22 | GRP1.8 | Exp 1 | 3 | 66.7 | 450.0 | 33.3 | 100.0 |
| 10 | GRP 2.23 | GRP1.8 | Exp 1 | 3 | 33.3 | 416.7 | 33.3 | 66.7 |
| 11 | GRP 2.24 | GRP1.8 | Exp 1 | 3 | 100 | 1600.0 | | 100.0 |
| 12 | GRP 2.3 | GRP1.8 | Exp 1 | 3 | 33.3 | 416.7 | 0.0 | 33.3 |
| 13 | GRP 2.6 | GRP1.8 | Exp 1 | 3 | 33.3 | 333.3 | 0.0 | 33.3 |
| 14 | GRP 2.7 | GRP1.8 | Exp 1 | 3 | 33.3 | 533.3 | 0.0 | 33.3 |
| 15 | GRP 2.9 | GRP1.8 | Exp 1 | 3 | 66.7 | 1066.7 | 0.0 | 66.7 |
| 16 | GRP 3.10 | GRP1.8 | Exp 1 | 3 | 33.3 | 416.7 | 0.0 | 33.3 |
| 17 | GRP 3.13 | GRP1.8 | Exp 1 | 3 | 100.0 | 1333.3 | | 100.0 |
| 18 | GRP 3.14 | GRP1.8 | Exp 1 | 3 | 66.7 | 750.0 | 0.0 | 66.7 |
| 19 | GRP 3.17 | GRP1.8 | Exp 1 | 3 | 100.0 | 1450.0 | | 100.0 |
| 20 | GRP 3.18 | GRP1.8 | Exp 1 | 3 | 66.7 | 633.3 | 0.0 | 66.7 |
| 21 | GRP 3.19 | GRP1.8 | Exp 1 | 3 | 33.3 | 500.0 | 0.0 | 33.3 |
| 22 | GRP 3.2 | GRP1.8 | Exp 1 | 3 | 66.7 | 683.3 | 0.0 | 66.7 |
| 23 | GRP 3.21 | GRP1.8 | Exp 1 | 3 | 66.7 | 866.7 | 0.0 | 66.7 |
| 24 | GRP 3.25 | GRP1.8 | Exp 1 | 3 | 33.3 | 533.3 | 0.0 | 33.3 |
| 25 | GRP 3.26 | GRP1.8 | Exp 1 | 3 | 33.3 | 333.3 | 0.0 | 33.3 |
| 26 | GRP 3.27 | GRP1.8 | Exp 1 | 3 | 100.0 | 1083.3 | | 100.0 |
| 27 | GRP 3.28 | GRP1.8 | Exp 1 | 3 | 66.7 | 516.7 | 0.0 | 66.7 |
| 28 | GRP 3.29 | GRP1.8 | Exp 1 | 3 | 100.0 | 466.7 | | 100.0 |
| 29 | GRP 3.32 | GRP1.8 | Exp 1 | 3 | 100.0 | 550.0 | | 100.0 |
| 30 | GRP 3.33 | GRP1.8 | Exp 1 | 3 | 100.0 | 1566.7 | | 100.0 |
| 31 | GRP 3.35 | GRP1.8 | Exp 1 | 3 | 0.0 | 0.0 | 33.3 | 33.3 |
| 32 | GRP 3.38 | GRP1.8 | Exp 1 | 3 | 100.0 | 1250.0 | | 100.0 |

| | | | | | | | | | |
|---|---|---|---|---|---|---|---|---|---|
| 33 | | GRP 3.39 | GRP1.8 | Exp 1 | 3 | 66.7 | 683.3 | 0.0 | 66.7 |
| 34 | | GRP 3.40 | GRP1.8 | Exp 1 | 3 | 66.7 | 1000.0 | 0.0 | 66.7 |
| 35 | | GRP 3.42 | GRP1.8 | Exp 1 | 3 | 100.0 | 1283.3 | | 100.0 |
| 36 | | GRP 3.45 | GRP1.8 | Exp 1 | 3 | 33.3 | 333.3 | 0.0 | 33.3 |
| 37 | | GRP 3.7 | GRP1.8 | Exp 1 | 3 | 66.7 | 633.3 | 0.0 | 66.7 |
| 38 | | GRP 3.9 | GRP1.8 | Exp 1 | 3 | 66.7 | 716.7 | 0.0 | 66.7 |
| 39 | | GRP 4.10 | GRP1.8 | Exp 1 | 3 | 33.3 | 416.7 | 0.0 | 33.3 |
| 40 | | GRP 4.11 | GRP1.8 | Exp 1 | 3 | 100.0 | 700.0 | | 100.0 |
| 41 | | GRP 4.13 | GRP1.8 | Exp 1 | 3 | 33.3 | 500.0 | 33.3 | 66.7 |
| 42 | | GRP 4.15 | GRP1.8 | Exp 1 | 3 | 33.3 | 533.3 | 0.0 | 33.3 |
| 43 | | GRP 4.16 | GRP1.8 | Exp 1 | 3 | 33.3 | 533.3 | 0.0 | 33.3 |
| 44 | | GRP 4.17 | GRP1.8 | Exp 1 | 3 | 66.7 | 1066.7 | 33.3 | 100.0 |
| 45 | | GRP 4.18 | GRP1.8 | Exp 1 | 3 | 66.7 | 950.0 | 33.3 | 100.0 |
| 46 | | GRP 4.19 | GRP1.8 | Exp 1 | 3 | 100.0 | 1400.0 | | 100.0 |
| 47 | | GRP 4.21 | GRP1.8 | Exp 1 | 3 | 100.0 | 1366.7 | | 100.0 |
| 48 | | GRP 4.28 | GRP1.8 | Exp 1 | 3 | 66.7 | 1066.7 | 0.0 | 66.7 |
| 49 | | GRP 4.5 | GRP1.8 | Exp 1 | 3 | 100.0 | 1366.7 | | 100.0 |
| 50 | | GRP 4.6 | GRP1.8 | Exp 1 | 3 | 66.7 | 950.0 | 0.0 | 66.7 |
| 51 | | GRP 4.8 | GRP1.8 | Exp 1 | 3 | 66.7 | 833.3 | 0.0 | 66.7 |
| 52 | | GRP 5.1 | GRP1.8 | Exp 1 | 3 | 100.0 | 1566.7 | | 100.0 |
| 53 | | GRP 5.10 | GRP1.8 | Exp 1 | 3 | 66.7 | 950.0 | 33.3 | 100.0 |
| 54 | | GRP 5.11 | GRP1.8 | Exp 1 | 3 | 100.0 | 1366.7 | | 100.0 |
| 55 | | GRP 5.15 | GRP1.8 | Exp 1 | 3 | 66.7 | 1033.3 | 0.0 | 66.7 |
| 56 | | GRP 5.16 | GRP1.8 | Exp 1 | 3 | 66.7 | 683.3 | 0.0 | 66.7 |
| 57 | | GRP 5.17 | GRP1.8 | Exp 1 | 3 | 33.3 | 183.3 | 33.3 | 66.7 |
| 58 | | GRP 5.18 | GRP1.8 | Exp 1 | 3 | 100.0 | 1083.3 | | 100.0 |
| 59 | | GRP 5.3 | GRP1.8 | Exp 1 | 3 | 66.7 | 1066.7 | 0.0 | 66.7 |
| 60 | | GRP 5.4 | GRP1.8 | Exp 1 | 3 | 100.0 | 1283.3 | | 100.0 |
| 61 | | GRP 5.5 | GRP1.8 | Exp 1 | 3 | 100.0 | 1050.0 | | 100.0 |
| 62 | | GRP 5.6 | GRP1.8 | Exp 1 | 3 | 100.0 | 1200.0 | | 100.0 |
| | 1 | II 10.4 | 35S | Exp 1 | 3 | 100.0 | 1216.7 | | 100.0 |
| | 2 | II 5.11 | 35S | Exp 1 | 3 | 100.0 | 1183.3 | | 100.0 |
| | 3 | II 7.13 | 35S | Exp 1 | 3 | 100.0 | 1216.7 | | 100.0 |
| | 4 | II 7.6 | 35S | Exp 1 | 3 | 66.7 | 950.0 | 0.0 | 66.7 |
| | 5 | II 7.9 | 35S | Exp 1 | 3 | 33.3 | 533.3 | 33.3 | 66.7 |
| | 6 | II 8.1 | 35S | Exp 1 | 3 | 100.0 | 1200.0 | | 100.0 |
| | 7 | II 8.5 | 35S | Exp 1 | 3 | 66.7 | 716.7 | 0.0 | 66.7 |
| | 8 | II 8.7 | 35S | Exp 1 | 3 | 100.0 | 1450.0 | | 100 |
| | 9 | II 9.3 | 35S | Exp 1 | 3 | 33.3 | 533.3 | 0.0 | 33.3 |
| | 10 | II G2 | 35S | Exp 1 | 3 | 100.0 | 1366.7 | | 100.0 |

| | | | | | | | | | |
|---|---|---|---|---|---|---|---|---|---|
| | 11 | II K | 35S | Exp 1 | 3 | 33.3 | 333.3 | 0.0 | 33.3 |
| | | Cruza 148 | Control | Exp 2 | 3 | 100.0 | 1716.7 | | 100.0 |
| | | Desiree | Control | Exp 2 | 3 | 100.0 | 1133.3 | | 100.0 |
| 63 | | GRP 4.23 | GRP1.8 | Exp 2 | 3 | 66.7 | 716.7 | 0.0 | 66.7 |
| 64 | | GRP 1.5 | GRP1.8 | Exp 2 | 3 | 0.0 | 0.0 | 0.0 | 0.0 |
| 65 | | GRP 3.11 | GRP1.8 | Exp 2 | 3 | 0.0 | 0.0 | 0.0 | 0.0 |
| 66 | | GRP 3.12 | GRP1.8 | Exp 2 | 3 | 100.0 | 1600.0 | | 100.0 |
| 67 | | GRP 3.15 | GRP1.8 | Exp 2 | 3 | 33.3 | 766.7 | 33.3 | 66.7 |
| 68 | | GRP 3.23 | GRP1.8 | Exp 2 | 3 | 66.7 | 1183.3 | 0.0 | 66.7 |
| 69 | | GRP 3.3 | GRP1.8 | Exp 2 | 3 | 66.7 | 716.7 | 33.3 | 100 |
| 70 | | GRP 3.30 | GRP1.8 | Exp 2 | 3 | 33.3 | 833.3 | 33.3 | 66.7 |
| 71 | | GRP 3.4 | GRP1.8 | Exp 2 | 3 | 100.0 | 1833.3 | | 100.0 |
| 72 | | GRP 4.20 | GRP1.8 | Exp 2 | 3 | 33.3 | 416.7 | 33.3 | 66.7 |
| 73 | | GRP 4.29 | GRP1.8 | Exp 2 | 3 | 100.0 | 2133.3 | | 100.0 |
| 74 | | GRP 5 | GRP1.8 | Exp 2 | 3 | 66.7 | 1600.0 | 0.0 | 66.7 |
| 75 | | GRP 5.20 | GRP1.8 | Exp 2 | 3 | 66.7 | 716.7 | 0.0 | 66.7 |
| 76 | | GRP 6.23 | GRP1.8 | Exp 2 | 3 | 0.0 | 0.0 | 33.3 | 33.3 |
| | 12 | II 3.35 | 35S | Exp 2 | 3 | 33.3 | 300.0 | 33.3 | 66.7 |
| | 13 | II 3.6 | 35S | Exp 2 | 3 | 66.7 | 1066.7 | 0.0 | 66.7 |
| | 14 | II 4.13 | 35S | Exp 2 | 3 | 33.3 | 766.7 | 0.0 | 33.3 |
| | 15 | II 4.16 | 35S | Exp 2 | 3 | 33.3 | 766.7 | 33.3 | 66.7 |
| | 16 | II 4.19 | 35S | Exp 2 | 3 | 100.0 | 1783.3 | | 100.0 |
| | 17 | II 4.33 | 35S | Exp 2 | 3 | 66.7 | 950.0 | 0.0 | 66.7 |
| | 18 | II 4.34 | 35S | Exp 2 | 3 | 66.7 | 1666.7 | 33.3 | 100.0 |
| | 19 | II 4.39 | 35S | Exp 2 | 3 | 66.7 | 1600.0 | 0.0 | 66.7 |
| | 20 | II 4.41 | 35S | Exp 2 | 3 | 66.7 | 1133.3 | 0.0 | 66.7 |
| | 21 | II 4.52 | 35S | Exp 2 | 3 | 100.0 | 2133.3 | | 100 |
| | 22 | II 4.60 | 35S | Exp 2 | 3 | 66.7 | 1533.3 | 0.0 | 66.7 |
| | 23 | II 5.2 | 35S | Exp 2 | 3 | 66.7 | 1666.7 | 0.0 | 66.7 |
| | 24 | II 6.11 | 35S | Exp 2 | 3 | 66.7 | 1016.7 | 0.0 | 66.7 |
| | 25 | II 6.12 | 35S | Exp 2 | 3 | 0.0 | 0.0 | 33.3 | 33.3 |
| | 26 | II 6.14 | 35S | Exp 2 | 3 | 66.7 | 1066.7 | 33.3 | 100.0 |
| | 27 | II 6.16 | 35S | Exp 2 | 3 | 0.0 | 0.0 | 0.0 | 0.0 |
| | 28 | II 6.20 | 35S | Exp 2 | 3 | 0.0 | 0.0 | 33.3 | 33.3 |
| | 29 | II 6.22 | 35S | Exp 2 | 3 | 33.3 | 766.7 | 0.0 | 33.3 |
| | 30 | II 6.24 | 35S | Exp 2 | 3 | 33.3 | 416.7 | 0.0 | 33.3 |
| | 31 | II 6.25 | 35S | Exp 2 | 3 | 100.0 | 2300.0 | 0.0 | 100.0 |
| | 32 | II 6.3 | 35S | Exp 2 | 3 | 0.0 | 0.0 | 0.0 | 0.0 |
| | 33 | II 6.6 | 35S | Exp 2 | 3 | 66.7 | 833.3 | 0.0 | 66.7 |

| | | | | | | | | | |
|---|---|---|---|---|---|---|---|---|---|
| | 34 | II 6.7 | 35S | Exp 2 | 3 | 0.0 | 0.0 | 33.3 | 33.3 |
| | 35 | II 6.8 | 35S | Exp 2 | 3 | 33.3 | 300.0 | 0.0 | 33.3 |
| | 36 | II 6.9 | 35S | Exp 2 | 3 | 66.7 | 1183.3 | 0.0 | 66.7 |
| | 37 | II H | 35S | Exp 2 | 3 | 33.3 | 650.0 | 33.3 | 66.7 |
| | 38 | II X | 35S | Exp 2 | 3 | 66.7 | 1600.0 | 33.3 | 100.0 |
| | | Cruza 148 | Control | Exp 3 | 3 | 0.0 | 0.0 | 33.3 | 33.3 |
| | | Desiree | Control | Exp 3 | 3 | 66.7 | 1533.3 | 0.0 | 66.7 |
| 77 | | GRP 11.1 | GRP1.8 | Exp 3 | 3 | 66.7 | 1533.3 | 33.3 | 100.0 |
| 78 | | GRP 11.2 | GRP1.8 | Exp 3 | 3 | 0.0 | 0.0 | 0.0 | 0.0 |
| 79 | | GRP 2.20 | GRP1.8 | Exp 3 | 3 | 100.0 | 1483.3 | | 100.0 |
| 80 | | GRP 2.8 | GRP1.8 | Exp 3 | 3 | 100.0 | 1483.3 | | 100.0 |
| 81 | | GRP 3.22 | GRP1.8 | Exp 3 | 3 | 100.0 | 2066.7 | | 100.0 |
| 82 | | GRP 4.33 | GRP1.8 | Exp 3 | 3 | 100.0 | 2416.7 | | 100.0 |
| 83 | | GRP 5.14 | GRP1.8 | Exp 3 | 3 | 33.3 | 533.3 | 0.0 | 33.3 |
| 84 | | GRP 5.2 | GRP1.8 | Exp 3 | 3 | 33.3 | 766.7 | 0.0 | 33.3 |
| 85 | | GRP 5.22 | GRP1.8 | Exp 3 | 3 | 66.7 | 1650 | 33.3 | 100.0 |
| 86 | | GRP 5.25 | GRP1.8 | Exp 3 | 3 | 100.0 | 2233.3 | | 100.0 |
| 87 | | GRP 5.26 | GRP1.8 | Exp 3 | 3 | 100.0 | 1950 | | 100.0 |
| 88 | | GRP 5.27 | GRP1.8 | Exp 3 | 3 | 100.0 | 1716.7 | | 100.0 |
| 89 | | GRP 5.28 | GRP1.8 | Exp 3 | 3 | 33.3 | 533.3 | 0.0 | 33.3 |
| 90 | | GRP 5.30 | GRP1.8 | Exp 3 | 3 | 100.0 | 1833.3 | | 100.0 |
| 91 | | GRP 5.31 | GRP1.8 | Exp 3 | 3 | 66.7 | 833.3 | 0.0 | 66.7 |
| 92 | | GRP 5.32 | GRP1.8 | Exp 3 | 3 | 66.7 | 1300 | 0.0 | 66.7 |
| 93 | | GRP 5.33 | GRP1.8 | Exp 3 | 3 | 100.0 | 1366.7 | | 100.0 |
| 94 | | GRP 5.34 | GRP1.8 | Exp 3 | 3 | 100.0 | 1483.3 | | 100.0 |
| 95 | | GRP 5.36 | GRP1.8 | Exp 3 | 3 | 100.0 | 1950.0 | | 100.0 |
| 96 | | GRP 5.37 | GRP1.8 | Exp 3 | 3 | 33.3 | 766.7 | 0.0 | 33.3 |
| 97 | | GRP 5.38 | GRP1.8 | Exp 3 | 3 | 100.0 | 2533.3 | | 100.0 |
| 98 | | GRP 5.39 | GRP1.8 | Exp 3 | 3 | 66.7 | 1116.7 | 0.0 | 66.7 |
| 99 | | GRP 5.40 | GRP1.8 | Exp 3 | 3 | 100.0 | 2416.7 | | 100.0 |
| 100 | | GRP 5.41 | GRP1.8 | Exp 3 | 3 | 66.7 | 1183.3 | 0.0 | 66.7 |
| 101 | | GRP 5.42 | GRP1.8 | Exp 3 | 3 | 100.0 | 2416.7 | | 100.0 |
| 102 | | GRP 5.44 | GRP1.8 | Exp 3 | 3 | 100.0 | 2183.3 | | 100.0 |
| 103 | | GRP 5.45 | GRP1.8 | Exp 3 | 3 | 100.0 | 2300.0 | | 100.0 |
| 104 | | GRP 5.50 | GRP1.8 | Exp 3 | 3 | 100.0 | 2066.7 | | 100.0 |
| 105 | | GRP 5.51 | GRP1.8 | Exp 3 | 3 | 100.0 | 2416.7 | | 100.0 |
| 106 | | GRP 5.52 | GRP1.8 | Exp 3 | 3 | 66.7 | 1416.7 | 33.3 | 100.0 |
| 107 | | GRP 5.53 | GRP1.8 | Exp 3 | 3 | 66.7 | 1533.3 | 33.3 | 100.0 |
| 108 | | GRP 5.56 | GRP1.8 | Exp 3 | 3 | 100.0 | 2533.3 | | 100.0 |

| | | | | | | | | |
|---|---|---|---|---|---|---|---|---|
| 109 | GRP 5.57 | GRP1.8 | Exp 3 | 3 | 66.7 | 1533.3 | 0.0 | 66.7 |
| 110 | GRP 5.60 | GRP1.8 | Exp 3 | 3 | 100.0 | 1716.7 | | 100.0 |
| 111 | GRP 5.9 | GRP1.8 | Exp 3 | 3 | 100.0 | 2533.3 | | 100.0 |
| 112 | GRP 6.1 | GRP1.8 | Exp 3 | 3 | 100.0 | 1950.0 | | 100.0 |
| 113 | GRP 6.10 | GRP1.8 | Exp 3 | 3 | 66.7 | 250.0 | 0.0 | 66.7 |
| 114 | GRP 6.11 | GRP1.8 | Exp 3 | 3 | 66.7 | 1300.0 | 33.3 | 100.0 |
| 115 | GRP 6.12 | GRP1.8 | Exp 3 | 3 | 33.3 | 533.3 | 0.0 | 33.3 |
| 116 | GRP 6.14 | GRP1.8 | Exp 3 | 3 | 66.7 | 1300.0 | 0.0 | 66.7 |
| 117 | GRP 6.18 | GRP1.8 | Exp 3 | 3 | 100.0 | 2066.7 | | 100.0 |
| 118 | GRP 6.19 | GRP1.8 | Exp 3 | 3 | 100.0 | 1250.0 | | 100.0 |
| 119 | GRP 6.2 | GRP1.8 | Exp 3 | 3 | 100.0 | 2416.7 | | 100.0 |
| 120 | GRP 6.22 | GRP1.8 | Exp 3 | 3 | 66.7 | 1650.0 | 33.3 | 100.0 |
| 121 | GRP 6.3 | GRP1.8 | Exp 3 | 3 | 100.0 | 2533.3 | | 100.0 |
| 122 | GRP 6.30 | GRP1.8 | Exp 3 | 3 | 100.0 | 2300.0 | | 100.0 |
| 123 | GRP 6.31 | GRP1.8 | Exp 3 | 3 | 100.0 | 2533.3 | | 100.0 |
| 124 | GRP 6.32 | GRP1.8 | Exp 3 | 3 | 100.0 | 1483.3 | | 100.0 |
| 125 | GRP 6.36 | GRP1.8 | Exp 3 | 3 | 66.7 | 833.3 | 33.3 | 100.0 |
| 126 | GRP 6.38 | GRP1.8 | Exp 3 | 3 | 33.3 | 650.0 | 33.3 | 66.7 |
| 127 | GRP 6.39 | GRP1.8 | Exp 3 | 3 | 66.7 | 1300.0 | 0.0 | 66.7 |
| 128 | GRP 6.42 | GRP1.8 | Exp 3 | 3 | 100.0 | 1950.0 | | 100.0 |
| 129 | GRP 6.43 | GRP1.8 | Exp 3 | 3 | 100.0 | 666.7 | | 100.0 |
| 130 | GRP 6.44 | GRP1.8 | Exp 3 | 3 | 33.3 | 933.3 | 66.7 | 100.0 |
| 131 | GRP 6.46 | GRP1.8 | Exp 3 | 3 | 100.0 | 1833.3 | | 100.0 |
| 132 | GRP 6.47 | GRP1.8 | Exp 3 | 3 | 66.7 | 1650.0 | 33.3 | 100.0 |
| 133 | GRP 6.48 | GRP1.8 | Exp 3 | 3 | 33.3 | 300.0 | 0.0 | 33.3 |
| 134 | GRP 6.49 | GRP1.8 | Exp 3 | 3 | 100.0 | 2066.7 | | 100.0 |
| 135 | GRP 6.5 | GRP1.8 | Exp 3 | 3 | 100.0 | 2533.3 | | 100.0 |
| 136 | GRP 6.51 | GRP1.8 | Exp 3 | 3 | 100.0 | 2183.3 | | 100.0 |
| 137 | GRP 6.52 | GRP1.8 | Exp 3 | 3 | 33.3 | 650.0 | 0 | 33.3 |
| 138 | GRP 6.53 | GRP1.8 | Exp 3 | 3 | 33.3 | 766.7 | 0 | 33.3 |
| 139 | GRP 6.55 | GRP1.8 | Exp 3 | 3 | 100.0 | 2066.7 | | 100.0 |
| 140 | GRP 6.56 | GRP1.8 | Exp 3 | 3 | 100.0 | 1950.0 | | 100.0 |
| 141 | GRP 6.57 | GRP1.8 | Exp 3 | 3 | 100.0 | 1950.0 | | 100.0 |
| 142 | GRP 6.9 | GRP1.8 | Exp 3 | 3 | 66.7 | 950 | 0 | 66.7 |
| 143 | GRP 7.1 | GRP1.8 | Exp 3 | 3 | 66.7 | 1066.7 | 33.3 | 100.0 |
| 144 | GRP 7.13 | GRP1.8 | Exp 3 | 3 | 100.0 | 2416.7 | | 100.0 |
| 145 | GRP 7.14 | GRP1.8 | Exp 3 | 3 | 100.0 | 2066.7 | | 100.0 |
| 146 | GRP 7.15 | GRP1.8 | Exp 3 | 3 | 33.3 | 66.7 | 0 | 33.3 |
| 147 | GRP 7.17 | GRP1.8 | Exp 3 | 3 | 100.0 | 1600.0 | | 100.0 |
| 148 | GRP 7.18 | GRP1.8 | Exp 3 | 3 | 33.3 | 533.3 | 0 | 33.3 |

| | | | | | | | | | |
|---|---|---|---|---|---|---|---|---|---|
| 149 | | GRP 7.19 | GRP1.8 | Exp 3 | 3 | 66.7 | 950.0 | 0 | 66.7 |
| 150 | | GRP 7.20 | GRP1.8 | Exp 3 | 3 | 100.0 | 2066.7 | | 100.0 |
| 151 | | GRP 7.21 | GRP1.8 | Exp 3 | 3 | 66.7 | 1533.3 | 0 | 66.7 |
| 152 | | GRP 7.27 | GRP1.8 | Exp 3 | 3 | 100.0 | 2183.3 | | 100.0 |
| 153 | | GRP 7.28 | GRP1.8 | Exp 3 | 3 | 0 | 0 | 33.3 | 33.3 |
| 154 | | GRP 7.3 | GRP1.8 | Exp 3 | 3 | 100.0 | 1833.3 | | 100.0 |
| 155 | | GRP 7.31 | GRP1.8 | Exp 3 | 3 | 100.0 | 2300.0 | | 100.0 |
| 156 | | GRP 7.32 | GRP1.8 | Exp 3 | 3 | 0.0 | 0.0 | 0.0 | 0.0 |
| 157 | | GRP 7.34 | GRP1.8 | Exp 3 | 3 | 66.7 | 1300.0 | 0.0 | 66.7 |
| 158 | | GRP 7.35 | GRP1.8 | Exp 3 | 3 | 100.0 | 2300.0 | | 100.0 |
| 159 | | GRP 7.36 | GRP1.8 | Exp 3 | 3 | 66.7 | 1533.3 | 0 | 66.7 |
| 160 | | GRP 7.37 | GRP1.8 | Exp 3 | 3 | 100.0 | 2300.0 | | 100.0 |
| 161 | | GRP 7.4 | GRP1.8 | Exp 3 | 3 | 100.0 | 2416.7 | | 100.0 |
| 162 | | GRP 7.5 | GRP1.8 | Exp 3 | 3 | 100.0 | 2300.0 | | 100.0 |
| 163 | | GRP 7.8 | GRP1.8 | Exp 3 | 3 | 100.0 | 2066.7 | | 100.0 |
| 164 | | GRP 8.1 | GRP1.8 | Exp 3 | 3 | 100.0 | 1716.7 | | 100.0 |
| 165 | | GRP 8.11 | GRP1.8 | Exp 3 | 3 | 66.7 | 133.3 | 33.3 | 100.0 |
| 166 | | GRP 8.12 | GRP1.8 | Exp 3 | 3 | 66.7 | 483.3 | 0.0 | 66.7 |
| 167 | | GRP 8.15 | GRP1.8 | Exp 3 | 3 | 0.0 | 0.0 | 33.3 | 33.3 |
| 168 | | GRP 8.17 | GRP1.8 | Exp 3 | 3 | 66.7 | 950.0 | 0.0 | 66.7 |
| 169 | | GRP 8.18 | GRP1.8 | Exp 3 | 3 | 66.7 | 1533.3 | 0.0 | 66.7 |
| 170 | | GRP 8.4 | GRP1.8 | Exp 3 | 3 | 66.7 | 1533.3 | 0.0 | 66.7 |
| 171 | | GRP 8.7 | GRP1.8 | Exp 3 | 3 | 100.0 | 1483.3 | | 100.0 |
| 172 | | GRP 8.8 | GRP1.8 | Exp 3 | 3 | 100.0 | 900.0 | | 100.0 |
| 173 | | GRP 9.2 | GRP1.8 | Exp 3 | 3 | 100.0 | 2300.0 | | 100.0 |
| 174 | | GRP 9.4 | GRP1.8 | Exp 3 | 3 | 33.3 | 533.3 | 0.0 | 33.3 |
| 175 | | GRP 9.5 | GRP1.8 | Exp 3 | 3 | 66.7 | 1183.3 | 0.0 | 66.7 |
| 176 | | GRP 9.6 | GRP1.8 | Exp 3 | 3 | 100 | 2066.7 | | 100.0 |
| 177 | | GRP 9.7 | GRP1.8 | Exp 3 | 3 | 0.0 | 0.0 | 0.0 | 0.0 |
| 178 | | GRP 9.8 | GRP1.8 | Exp 3 | 3 | 33.3 | 883.3 | 0.0 | 33.3 |
| | | Cruza 148 | Control | Exp 4 | 3 | 66.7 | 1383.3 | 33.3 | 100.0 |
| | | Desiree | Control | Exp 4 | 3 | 100.0 | 2016.7 | | 100.0 |
| 179 | | GRP 8.3 | GRP1.8 | Exp 4 | 3 | 33.3 | 633.3 | 0.0 | 33.3 |
| 180 | | GRP 8.6 | GRP1.8 | Exp 4 | 3 | 66.7 | 1616.7 | 0.0 | 66.7 |
| | 39 | II 6.23 | 35S | Exp 4 | 3 | 100.0 | 2600.0 | | 100.0 |
| | 40 | II 10.1 | 35S | Exp 4 | 3 | 100.0 | 2600.0 | | 100.0 |
| | 41 | II 10.11 | 35S | Exp 4 | 3 | 100.0 | 1900.0 | | 100.0 |
| | 42 | II 10.12 | 35S | Exp 4 | 3 | 66.7 | 1383.3 | 0.0 | 66.7 |
| | 43 | II 10.13 | 35S | Exp 4 | 3 | 100.0 | 2366.7 | | 100.0 |

| | | | | | | | | |
|---|---|---|---|---|---|---|---|---|
| 44 | II 10.14 | 35S | Exp 4 | 3 | 66.7 | 1733.3 | 0.0 | 66.7 |
| 45 | II 10.15 | 35S | Exp 4 | 3 | 66.7 | 1800 | 0.0 | 66.7 |
| 46 | II 10.16 | 35S | Exp 4 | 3 | 100.0 | 2366.7 | | 100.0 |
| 47 | II 10.19 | 35S | Exp 4 | 3 | 100.0 | 2550.0 | | 100.0 |
| 48 | II 10.20 | 35S | Exp 4 | 3 | 66.7 | 916.7 | 0.0 | 66.7 |
| 49 | II 10.21 | 35S | Exp 4 | 3 | 100.0 | 2483.3 | | 100.0 |
| 50 | II 10.23 | 35S | Exp 4 | 3 | 100.0 | 2483.3 | | 100.0 |
| 51 | II 10.24 | 35S | Exp 4 | 3 | 66.7 | 1266.7 | 33.3 | 100.0 |
| 52 | II 10.25 | 35S | Exp 4 | 3 | 100.0 | 1900.0 | | 100.0 |
| 53 | II 10.26 | 35S | Exp 4 | 3 | 100.0 | 2316.7 | | 100.0 |
| 54 | II 10.29 | 35S | Exp 4 | 3 | 66.7 | 1033.3 | 0.0 | 66.7 |
| 55 | II 10.3 | 35S | Exp 4 | 3 | 0.0 | 0.0 | 66.7 | 66.7 |
| 56 | II 10.33 | 35S | Exp 4 | 3 | 100.0 | 2600.0 | | 100.0 |
| 57 | II 10.5 | 35S | Exp 4 | 3 | 66.7 | 1800.0 | 0.0 | 66.7 |
| 58 | II 10.7 | 35S | Exp 4 | 3 | 100.0 | 2600.0 | | 100.0 |
| 59 | II 10.8 | 35S | Exp 4 | 3 | 100.0 | 2550.0 | | 100.0 |
| 60 | II 11.12 | 35S | Exp 4 | 3 | 100.0 | 1550.0 | | 100.0 |
| 61 | II 11.14 | 35S | Exp 4 | 3 | 66.7 | 1733.3 | 0.0 | 66.7 |
| 62 | II 11.15 | 35S | Exp 4 | 3 | 100.0 | 2133.3 | | 100.0 |
| 63 | II 11.16 | 35S | Exp 4 | 3 | 100.0 | 2250.0 | | 100.0 |
| 64 | II 11.17 | 35S | Exp 4 | 3 | 100.0 | 1666.7 | | 100.0 |
| 65 | II 11.18 | 35S | Exp 4 | 3 | 100.0 | 2733.3 | | 100.0 |
| 66 | II 11.19 | 35S | Exp 4 | 3 | 100.0 | 1900.0 | | 100.0 |
| 67 | II 11.2 | 35S | Exp 4 | 3 | 66.7 | 916.7 | 0.0 | 66.7 |
| 68 | II 11.21 | 35S | Exp 4 | 3 | 100.0 | 2433.3 | | 100.0 |
| 69 | II 11.22 | 35S | Exp 4 | 3 | 33.3 | 633.3 | 0.0 | 33.3 |
| 70 | II 11.4 | 35S | Exp 4 | 3 | 100.0 | 1900.0 | | 100.0 |
| 71 | II 11.5 | 35S | Exp 4 | 3 | 33.3 | 866.7 | 33.3 | 66.7 |
| 72 | II 11.7 | 35S | Exp 4 | 3 | 100.0 | 2016.7 | | 100.0 |
| 73 | II 12.11 | 35S | Exp 4 | 3 | 100.0 | 1783.3 | | 100.0 |
| 74 | II 12.2 | 35S | Exp 4 | 3 | 100.0 | 2016.7 | | 100.0 |
| 75 | II 12.3 | 35S | Exp 4 | 3 | 0.0 | 0.0 | 0.0 | 0.0 |
| 76 | II 12.4 | 35S | Exp 4 | 3 | 100.0 | 2483.3 | | 100.0 |
| 77 | II 12.7 | 35S | Exp 4 | 3 | 100.0 | 2483.3 | | 100.0 |
| 78 | II 13.2 | 35S | Exp 4 | 3 | 66.7 | 1150 | 0.0 | 66.7 |
| 79 | II 13.4 | 35S | Exp 4 | 3 | 100.0 | 2666.7 | | 100.0 |
| 80 | II 14.1 | 35S | Exp 4 | 3 | 100.0 | 1900.0 | | 100.0 |
| 81 | II 14.2 | 35S | Exp 4 | 3 | 100.0 | 2250.0 | | 100.0 |
| 82 | II 14.3 | 35S | Exp 4 | 3 | 33.3 | 633.3 | 0.0 | 33.3 |
| 83 | II 14.4 | 35S | Exp 4 | 3 | 66.7 | 1033.3 | 0.0 | 66.7 |

| | | | | | | | | |
|---|---|---|---|---|---|---|---|---|
| 84 | II 14.5 | 35S | Exp 4 | 3 | 100.0 | 2250.0 | 0.0 | 100.0 |
| 85 | II 16.1 | 35S | Exp 4 | 3 | 66.7 | 1500.0 | 0.0 | 66.7 |
| 86 | II 17.2 | 35S | Exp 4 | 3 | 100.0 | 2366.7 | 0.0 | 100.0 |
| 87 | II 2.10 | 35S | Exp 4 | 3 | 66.7 | 1266.7 | 0.0 | 66.7 |
| 88 | II 3.2 | 35S | Exp 4 | 3 | 100.0 | 2366.7 | | 100.0 |
| 89 | II 3.20 | 35S | Exp 4 | 3 | 100.0 | 2016.7 | | 100.0 |
| 90 | II 3.22 | 35S | Exp 4 | 3 | 100.0 | 2483.3 | | 100.0 |
| 91 | II 3.23 | 35S | Exp 4 | 3 | 100.0 | 2366.7 | | 100.0 |
| 92 | II 3.26 | 35S | Exp 4 | 3 | 100.0 | 2016.7 | | 100.0 |
| 93 | II 3.28 | 35S | Exp 4 | 3 | 66.7 | 916.7 | 0.0 | 66.7 |
| 94 | II 3.32 | 35S | Exp 4 | 3 | 33.3 | 166.7 | 0.0 | 33.3 |
| 95 | II 3.4 | 35S | Exp 4 | 3 | 100.0 | 2016.7 | | 100.0 |
| 96 | II 3.41 | 35S | Exp 4 | 3 | 100.0 | 1783.3 | | 100.0 |
| 97 | II 4.10 | 35S | Exp 4 | 3 | 66.7 | 1500.0 | 0.0 | 66.7 |
| 98 | II 4.11 | 35S | Exp 4 | 3 | 66.7 | 1683.3 | 0.0 | 66.7 |
| 99 | II 4.14 | 35S | Exp 4 | 3 | 100.0 | 2250.0 | | 100.0 |
| 100 | II 4.15 | 35S | Exp 4 | 3 | 66.7 | 1616.7 | 0.0 | 66.7 |
| 101 | II 4.23 | 35S | Exp 4 | 3 | 66.7 | 1266.7 | 33.3 | 100.0 |
| 102 | II 4.29 | 35S | Exp 4 | 3 | 100.0 | 2366.7 | | 100.0 |
| 103 | II 4.32 | 35S | Exp 4 | 3 | 66.7 | 1266.7 | 0.0 | 66.7 |
| 104 | II 4.43 | 35S | Exp 4 | 3 | 100.0 | 1666.7 | | 100.0 |
| 105 | II 4.45 | 35S | Exp 4 | 3 | 66.7 | 1733.3 | 33.3 | 100.0 |
| 106 | II 4.5 | 35S | Exp 4 | 3 | 33.3 | 866.7 | 0.0 | 33.3 |
| 107 | II 4.53 | 35S | Exp 4 | 3 | 66.7 | 1733.3 | 0.0 | 66.7 |
| 108 | II 4.7 | 35S | Exp 4 | 3 | 100.0 | 2666.7 | | 100.0 |
| 109 | II 5.12 | 35S | Exp 4 | 3 | 66.7 | 1800.0 | 33.3 | 100.0 |
| 110 | II 5.18 | 35S | Exp 4 | 3 | 100.0 | 2133.3 | | 100.0 |
| 111 | II 5.22 | 35S | Exp 4 | 3 | 66.7 | 1333.3 | 0.0 | 66.7 |
| 112 | II 5.3 | 35S | Exp 4 | 3 | 100.0 | 2600.0 | | 100.0 |
| 113 | II 5.5 | 35S | Exp 4 | 3 | 100.0 | 2483.3 | | 100.0 |
| 114 | II 5.8 | 35S | Exp 4 | 3 | 66.7 | 800.0 | 0.0 | 66.7 |
| 115 | II 5.9 | 35S | Exp 4 | 3 | 100.0 | 2366.7 | | 100.0 |
| 116 | II 6.28 | 35S | Exp 4 | 3 | 100.0 | 2016.7 | | 100.0 |
| 117 | II 6.30 | 35S | Exp 4 | 3 | 100.0 | 2200.0 | | 100.0 |
| 118 | II 6.31 | 35S | Exp 4 | 3 | 100.0 | 2550.0 | | 100.0 |
| 119 | II 6.33 | 35S | Exp 4 | 3 | 66.7 | 1383.3 | 0.0 | 66.7 |
| 120 | II 6.35 | 35S | Exp 4 | 3 | 100.0 | 2600.0 | | 100.0 |
| 121 | II 6.36 | 35S | Exp 4 | 3 | 100.0 | 1666.7 | | 100.0 |
| 122 | II 6.37 | 35S | Exp 4 | 3 | 100.0 | 2433.3 | | 100.0 |
| 123 | II 6.40 | 35S | Exp 4 | 3 | 100.0 | 1550.0 | | 100.0 |

| | | | | | | | | | |
|---|---|---|---|---|---|---|---|---|---|
| | 124 | II 6.41 | 35S | Exp 4 | 3 | 100.0 | 2383.3 | | 100.0 |
| | 125 | II 6.42 | 35S | Exp 4 | 3 | 100.0 | 2550.0 | | 100.0 |
| | 126 | II 6.44 | 35S | Exp 4 | 3 | 100.0 | 1433.3 | | 100.0 |
| | 127 | II 6.45 | 35S | Exp 4 | 3 | 100.0 | 2133.3 | | 100.0 |
| | 128 | II 6.46 | 35S | Exp 4 | 3 | 66.7 | 1500.0 | 0.0 | 66.7 |
| | 129 | II 6.47 | 35S | Exp 4 | 3 | 66.7 | 1266.7 | 0.0 | 66.7 |
| | 130 | II 6.48 | 35S | Exp 4 | 3 | 33.3 | 166.7 | 0.0 | 33.3 |
| | 131 | II 6.49 | 35S | Exp 4 | 3 | 100.0 | 2383.3 | | 100.0 |
| | 132 | II 6.51 | 35S | Exp 4 | 3 | 100.0 | 1783.3 | | 100.0 |
| | 133 | II 6.52 | 35S | Exp 4 | 3 | 100.0 | 1433.3 | | 100.0 |
| | 134 | II 6.55 | 35S | Exp 4 | 3 | 100.0 | 2366.7 | | 100.0 |
| | 135 | II 6.56 | 35S | Exp 4 | 3 | 100.0 | 2250.0 | | 100.0 |
| | 136 | II 6.59 | 35S | Exp 4 | 3 | 66.7 | 1150.0 | 0.0 | 66.7 |
| | 137 | II 6.62 | 35S | Exp 4 | 3 | 100.0 | 2433.3 | | 100.0 |
| | 138 | II 6.64 | 35S | Exp 4 | 3 | 100.0 | 2500.0 | | 100.0 |
| | 139 | II 6.67 | 35S | Exp 4 | 3 | 100.0 | 2666.7 | | 100.0 |
| | 140 | II 6.70 | 35S | Exp 4 | 3 | 66.7 | 1733.3 | 0.0 | 66.7 |
| | 141 | II I | 35S | Exp 4 | 3 | 100.0 | 2083.3 | | 100.0 |
| | 142 | II L | 35S | Exp 4 | 3 | 100.0 | 2016.7 | | 100.0 |
| | 143 | II TE | 35S | Exp 4 | 3 | 100.0 | 2366.7 | | 100.0 |
| | | Cruza 148 | Control | Exp 5 | 3 | 0.0 | 0.0 | 0.0 | 0.0 |
| | | Desiree | Control | Exp 5 | 3 | 66.7 | 1066.7 | 0.0 | 66.7 |
| 181 | | GRP 12 | GRP1.8 | Exp 5 | 3 | 66.7 | 1466.7 | 0.0 | 66.7 |
| 182 | | GRP 2.15 | GRP1.8 | Exp 5 | 3 | 66.7 | 716.7 | 0.0 | 66.7 |
| 183 | | GRP 2.16 | GRP1.8 | Exp 5 | 3 | 33.3 | 416.7 | 0.0 | 33.3 |
| 184 | | GRP 5.46 | GRP1.8 | Exp 5 | 3 | 33.3 | 883.3 | 0.0 | 33.3 |
| 185 | | GRP 6.24 | GRP1.8 | Exp 5 | 3 | 66.7 | 1816.7 | 0.0 | 66.7 |
| 186 | | GRP 6.25 | GRP1.8 | Exp 5 | 3 | 33.3 | 933.3 | 0.0 | 33.3 |
| 187 | | GRP 6.33 | GRP1.8 | Exp 5 | 3 | 100.0 | 1483.3 | | 100.0 |
| 188 | | GRP 6.34 | GRP1.8 | Exp 5 | 3 | 100.0 | 1366.7 | | 100.0 |
| 189 | | GRP 6.35 | GRP1.8 | Exp 5 | 3 | 33.3 | 300.0 | 0.0 | 33.3 |
| 190 | | GRP 6.45 | GRP1.8 | Exp 5 | 3 | 66.7 | 1416.7 | 0.0 | 66.7 |
| 191 | | GRP 6.54 | GRP1.8 | Exp 5 | 3 | 0.0 | 0.0 | 0.0 | 0.0 |
| 192 | | GRP 6.58 | GRP1.8 | Exp 5 | 3 | 33.3 | 66.7 | 0.0 | 33.3 |
| 193 | | GRP 7.10 | GRP1.8 | Exp 5 | 3 | 33.3 | 533.3 | 0.0 | 33.3 |
| 194 | | GRP 7.12 | GRP1.8 | Exp 5 | 3 | 66.7 | 950.0 | 0.0 | 66.7 |
| 195 | | GRP 7.16 | GRP1.8 | Exp 5 | 3 | 66.7 | 1533.3 | 0.0 | 66.7 |
| 196 | | GRP 7.22 | GRP1.8 | Exp 5 | 3 | 33.3 | 883.3 | 0.0 | 33.3 |
| 197 | | GRP 8.10 | GRP1.8 | Exp 5 | 3 | 0.0 | 0.0 | 0.0 | 0.0 |

| | | | | | | | | |
|---|---|---|---|---|---|---|---|---|
| 144 | II 10.10 | 35S | Exp 5 | 3 | 100.0 | 1600.0 | | 100.0 |
| 145 | II 10.18 | 35S | Exp 5 | 3 | 0.0 | 0.0 | 33.3 | 33.3 |
| 146 | II 10.22 | 35S | Exp 5 | 3 | 66.7 | 1183.3 | 0.0 | 66.7 |
| 147 | II 10.31 | 35S | Exp 5 | 3 | 33.3 | 66.7 | 0.0 | 33.3 |
| 148 | II 11.3 | 35S | Exp 5 | 3 | 0.0 | 0.0 | 0.0 | 0.0 |
| 149 | II 11.6 | 35S | Exp 5 | 3 | 33.3 | 933.3 | 0.0 | 33.3 |
| 150 | II 12.5 | 35S | Exp 5 | 3 | 100.0 | 1016.7 | | 100.0 |
| 151 | II 13.3 | 35S | Exp 5 | 3 | 66.7 | 1183.3 | 0.0 | 66.7 |
| 152 | II 15.1 | 35S | Exp 5 | 3 | 66.7 | 1233.3 | 0.0 | 66.7 |
| 153 | II 4.20 | 35S | Exp 5 | 3 | 100.0 | 2233.3 | | 100.0 |
| 154 | II 4.36 | 35S | Exp 5 | 3 | 33.3 | 650.0 | 33.3 | 66.7 |
| 155 | II 4.44 | 35S | Exp 5 | 3 | 0.0 | 0.0 | 0.0 | 0.0 |
| 156 | II 5.17 | 35S | Exp 5 | 3 | 66.7 | 1466.7 | 0.0 | 66.7 |
| 157 | II 5.19 | 35S | Exp 5 | 3 | 0.0 | 0.0 | 33.3 | 33.3 |
| 158 | II 5.21 | 35S | Exp 5 | 3 | 33.3 | 933.3 | 33.3 | 66.7 |
| 159 | II 5.24 | 35S | Exp 5 | 3 | 33.3 | 300.0 | 66.7 | 100.0 |
| 160 | II 5.4 | 35S | Exp 5 | 3 | 66.7 | 950.0 | 0.0 | 66.7 |
| 161 | II 6.29 | 35S | Exp 5 | 3 | 0.0 | 0.0 | 0.0 | 0.0 |
| 162 | II Wa | 35S | Exp 5 | 3 | 66.7 | 1866.7 | 0.0 | 66.7 |
| 163 | II WB | 35S | Exp 5 | 3 | 66.7 | 1066.7 | 0.0 | 66.7 |

* Area under disease progress curve

**Transgenic events with 100% wilting were not tested for latent infection
