## Supplementary material for "Expression of the β*hpmeh* gene in transgenic events of the potato variety Desiree increases resistance to bacterial wilt caused by *Ralstonia solanacearum*": TableS2

**Table S2**: Statistical analysis of phenotypic results (Exp 6,7,9,10: Scott & Knott, and Exp 8,11-12: Kruskal-Wallis analysis method). Means with the same letter are not significantly different at the 5% level (Scott & Knott, 1974). Ranks with the same letter are not significantly different at the 5% level (De Mendiburu, 2012).

| Event | Promoter | Experiment (Exp) | Wilting (%) | mean rank | AUDPC* | mean rank | Latent Infection (%)** | Infected plants (%) | mean rank |
|---|---|---|---|---|---|---|---|---|---|
| Cruza 148 | control | Exp 6 | 6.7 | **c** | 100.0 | **c** | 20.0 | 26.7 | **c** |
| GRP 3.11 | GRP1.8 | Exp 6 | 6.7 | **c** | 113.3 | **c** | 6.7 | 13.3 | **c** |
| II 4.16 | 35S | Exp 6 | 40.0 | b | 576.7 | **c** | 13.3 | 53.3 | b |
| GRP 2.6 | GRP1.8 | Exp 6 | 66.7 | a | 1046.7 | b | 0.0 | 66.7 | a |
| GRP 3.19 | GRP1.8 | Exp 6 | 86.7 | a | 1273.3 | b | 0.0 | 86.7 | a |
| GRP 3.35 | GRP1.8 | Exp 6 | 73.3 | a | 1346.7 | b | 6.7 | 80.0 | a |
| II 6.3 | 35S | Exp 6 | 93.3 | a | 1426.7 | b | 0.0 | 93.3 | a |
| II 6.20 | 35S | Exp 6 | 86.7 | a | 1490.0 | b | 0.0 | 86.7 | a |
| Desiree | control | Exp 6 | 93.3 | a | 1523.3 | b | 0.0 | 93.3 | a |
| GRP 2.3 | GRP1.8 | Exp 6 | 86.7 | a | 1563.3 | b | 0.0 | 86.7 | a |
| GRP 3.10 | GRP1.8 | Exp 6 | 93.3 | a | 1603.3 | b | 0.0 | 93.3 | a |
| II 4.13 | 35S | Exp 6 | 86.7 | a | 1663.3 | b | 0.0 | 86.7 | a |
| GRP 2.24 | GRP1.8 | Exp 6 | 93.3 | a | 1676.7 | b | 0.0 | 93.3 | a |
| GRP 4.17 | GRP1.8 | Exp 6 | 86.7 | a | 1723.3 | a | 6.7 | 93.3 | a |
| GRP 3.26 | GRP1.8 | Exp 6 | 93.3 | a | 1776.7 | a | 0.0 | 93.3 | a |
| II 6.8 | 35S | Exp 6 | 100.0 | a | 1873.3 | a | | 100.0 | a |
| II 6.7 | 35S | Exp 6 | 93.3 | a | 1883.3 | a | 0.0 | 93.3 | a |
| II 6.12 | 35S | Exp 6 | 100.0 | a | 1913.3 | a | | 100.0 | a |
| II 6.22 | 35S | Exp 6 | 100.0 | a | 1913.3 | a | | 100.0 | a |
| II 6.16 | 35S | Exp 6 | 100.0 | a | 1920.0 | a | | 100.0 | a |
| GRP 1.5 | GRP1.8 | Exp 6 | 100.0 | a | 1993.3 | a | | 100.0 | a |
| GRP 2.18 | GRP1.8 | Exp 6 | 100.0 | a | 2010.0 | a | | 100.0 | a |
| GRP 3.45 | GRP1.8 | Exp 6 | 100.0 | a | 2076.7 | a | | 100.0 | a |

| | | | | | | | | | | | | |
|---|---|---|---|---|---|---|---|---|---|---|---|---|
| GRP 4.10 | GRP1.8 | Exp 6 | 100.0 | | a | 2096.7 | | a | | 100.0 | | a |
| II 4.16 | 35S | Exp 7 | 11.1 | | a | 133.3 | | a | 11.1 | 22.2 | | **b** |
| Cruza 148 | control | Exp 7 | 20.0 | | a | 263.3 | | a | 0.0 | 20.0 | | **b** |
| GRP 3.11 | GRP1.8 | Exp 7 | 26.7 | | a | 320.0 | | a | 6.7 | 33.3 | | **b** |
| GRP 3.35 | GRP1.8 | Exp 7 | 26.7 | | a | 390.0 | | a | 0.0 | 26.7 | | **b** |
| GRP 3.10 | GRP1.8 | Exp 7 | 33.3 | | a | 470.0 | | a | 0.0 | 33.3 | | **b** |
| GRP 4.17 | GRP1.8 | Exp 7 | 26.7 | | a | 473.3 | | a | 13.3 | 40.0 | | **b** |
| II 6.3 | 35S | Exp 7 | 33.3 | | a | 540.0 | | a | 6.7 | 40.0 | | **b** |
| GRP 2.6 | GRP1.8 | Exp 7 | 46.7 | | a | 560.0 | | a | 20.0 | 66.7 | | a |
| GRP 2.3 | GRP1.8 | Exp 7 | 40.0 | | a | 573.3 | | a | 0.0 | 40.0 | | **b** |
| II 6.22 | 35S | Exp 7 | 46.7 | | a | 606.7 | | a | 0.0 | 46.7 | | a |
| GRP 3.45 | GRP1.8 | Exp 7 | 40.0 | | a | 610.0 | | a | 0.0 | 40.0 | | **b** |
| II 6.8 | 35S | Exp 7 | 40.0 | | a | 633.3 | | a | 13.3 | 53.3 | | a |
| GRP 2.18 | GRP1.8 | Exp 7 | 46.7 | | a | 700.0 | | a | 0.0 | 46.7 | | a |
| II 6.16 | 35S | Exp 7 | 53.3 | | a | 746.7 | | a | 20.0 | 73.3 | | a |
| II 4.13 | 35S | Exp 7 | 40.0 | | a | 760.0 | | a | 0.0 | 40.0 | | **b** |
| GRP 3.26 | GRP1.8 | Exp 7 | 46.7 | | a | 770.0 | | a | 0.0 | 46.7 | | a |
| II 6.20 | 35S | Exp 7 | 40.0 | | a | 830.0 | | a | 13.3 | 53.3 | | a |
| II 6.12 | 35S | Exp 7 | 66.7 | | a | 846.7 | | a | 0.0 | 66.7 | | a |
| Desiree | control | Exp 7 | 60.0 | | a | 976.7 | | a | 0.0 | 60.0 | | a |
| GRP 1.5 | GRP1.8 | Exp 7 | 66.7 | | a | 1033.3 | | a | 13.3 | 80.0 | | a |
| GRP 2.24 | GRP1.8 | Exp 7 | 53.3 | | a | 1036.7 | | a | 6.7 | 60.0 | | a |
| GRP 4.10 | GRP1.8 | Exp 7 | 60.0 | | a | 1140.0 | | a | 6.7 | 66.7 | | a |
| GRP 3.19 | GRP1.8 | Exp 7 | 60.0 | | a | 1200.0 | | a | 13.3 | 73.3 | | a |
| II 6.7 | 35S | Exp 7 | 66.7 | | a | 1303.3 | | a | 0.0 | 66.7 | | a |
| GRP 3.11 | GRP1.8 | Exp 8 | 6.7 | 2.0 | **d** | 60.0 | 2.0 | **f** | 0.0 | 6.7 | 2.0 | **d** |
| II 4.16 | 35S | Exp 8 | 73.3 | 30.2 | abc | 730.0 | 9.8 | **ef** | 7.1 | 80.0 | 36.3 | abc |
| II 4.13 | 35S | Exp 8 | 80.0 | 33.3 | abc | 1124.4 | 18.2 | **def** | 0.0 | 80.0 | 30.2 | abc |

| | | | | | | | | | | | | |
|---|---|---|---|---|---|---|---|---|---|---|---|---|
| GRP 2.18 | GRP1.8 | Exp 8 | 60.0 | 17.0 | cd | 936.7 | 20.5 | **def** | 7.1 | 66.7 | 23.2 | bcd |
| GRP 2.6 | GRP1.8 | Exp 8 | 66.7 | 25.0 | bcd | 1123.3 | 22.5 | **cdef** | 0.0 | 66.7 | 23.2 | bcd |
| GRP 2.3 | GRP1.8 | Exp 8 | 60.0 | 17.0 | cd | 1100.0 | 22.8 | **bcdef** | 7.1 | 66.7 | 17.0 | cd |
| GRP 2.24 | GRP1.8 | Exp 8 | 80.0 | 33.3 | abc | 1151.7 | 24.3 | **bcdef** | 0.0 | 80.0 | 30.2 | abc |
| GRP 1.5 | GRP1.8 | Exp 8 | 66.7 | 20.2 | cd | 1230.0 | 24.7 | **bcdef** | 7.1 | 73.3 | 21.5 | bcd |
| II 6.7 | 35S | Exp 8 | 80.0 | 30.5 | abc | 1396.7 | 33.5 | abcde | 0.0 | 80.0 | 26.0 | bcd |
| Desiree | control | Exp 8 | 80.0 | 33.3 | abc | 1536.7 | 36.5 | abcde | 0.0 | 80.0 | 30.2 | abc |
| II 6.20 | 35S | Exp 8 | 70.0 | 25.8 | bcd | 1529.2 | 36.7 | abcde | 7.1 | 76.7 | 28.3 | abc |
| II 6.3 | 35S | Exp 8 | 86.7 | 41.3 | abc | 1443.3 | 37.7 | abcd | 7.1 | 93.3 | 43.3 | ab |
| II 6.16 | 35S | Exp 8 | 86.7 | 38.5 | abc | 1480.0 | 38.2 | abcd | 7.1 | 93.3 | 43.3 | ab |
| GRP 4.10 | GRP1.8 | Exp 8 | 80.0 | 33.3 | abc | 1449.2 | 38.2 | abcd | 14.3 | 93.3 | 43.3 | ab |
| GRP 3.26 | GRP1.8 | Exp 8 | 93.3 | 46.5 | ab | 1493.3 | 38.5 | abcd | 7.1 | 100.0 | 52.0 | a |
| II 6.12 | 35S | Exp 8 | 86.7 | 38.5 | abc | 1620.0 | 42.0 | abcd | 0.0 | 86.7 | 34.7 | abc |
| GRP 4.17 | GRP1.8 | Exp 8 | 100.0 | 54.5 | a | 1600.0 | 44.2 | abcd | | 100.0 | 52.0 | a |
| GRP 3.19 | GRP1.8 | Exp 8 | 73.3 | 28.2 | bcd | 1686.7 | 44.5 | abcd | 0.0 | 73.3 | 25.7 | bcd |
| GRP 3.35 | GRP1.8 | Exp 8 | 100.0 | 54.5 | a | 1734.2 | 49.0 | abc | | 100.0 | 52.0 | a |
| GRP 3.10 | GRP1.8 | Exp 8 | 93.3 | 46.5 | ab | 1750.0 | 50.0 | ab | 0.0 | 93.3 | 43.3 | ab |
| II 6.22 | 35S | Exp 8 | 100.0 | 54.5 | a | 1773.3 | 53.3 | a | | 100.0 | 52.0 | a |
| II 6.8 | 35S | Exp 8 | 100.0 | 54.5 | a | 1996.7 | 58.2 | a | | 100.0 | 52.0 | a |
| GRP 3.45 | GRP1.8 | Exp 8 | 93.3 | 46.5 | ab | 2030.0 | 59.8 | a | 0.0 | 93.3 | 43.3 | ab |
| GRP 7.28 | GRP1.8 | Exp 9 | 28.3 | | **b** | 336.7 | | **b** | 0.0 | 28.3 | | **b** |
| GRP 7.15 | GRP1.8 | Exp 9 | 40.0 | | **b** | 570.0 | | **b** | 0.0 | 40.0 | | **b** |
| Cruza 148 | control | Exp 9 | 46.7 | | **b** | 606.7 | | **b** | 0.0 | 46.7 | | **b** |
| GRP 9.7 | GRP1.8 | Exp 9 | 53.3 | | **b** | 690.0 | | **b** | 0.0 | 53.3 | | **b** |
| GRP 6.48 | GRP1.8 | Exp 9 | 46.7 | | **b** | 826.7 | | **b** | 0.0 | 46.7 | | **b** |
| GRP 6.12 | GRP1.8 | Exp 9 | 73.3 | | a | 870.0 | | **b** | 6.7 | 80.0 | | a |
| GRP 5.14 | GRP1.8 | Exp 9 | 63.3 | | a | 972.5 | | **b** | 0.0 | 63.3 | | **b** |
| GRP 6.52 | GRP1.8 | Exp 9 | 73.3 | | a | 1243.3 | | a | 0.0 | 73.3 | | a |

| | | | | | | | | | |
|---|---|---|---|---|---|---|---|---|---|
| GRP 7.32 | GRP1.8 | Exp 9 | 66.7 | a | 1253.3 | a | 0.0 | 66.7 | a |
| GRP 11.2 | GRP1.8 | Exp 9 | 73.3 | a | 1500.0 | a | 0.0 | 73.3 | a |
| GRP 5.2 | GRP1.8 | Exp 9 | 80.0 | a | 1560.0 | a | 0.0 | 80.0 | a |
| GRP 6.53 | GRP1.8 | Exp 9 | 80.0 | a | 1583.3 | a | 6.7 | 86.7 | a |
| Desiree | control | Exp 9 | 80.0 | a | 1700.0 | a | 6.7 | 86.7 | a |
| GRP 9.8 | GRP1.8 | Exp 9 | 86.7 | a | 1746.7 | a | 6.7 | 93.3 | a |
| GRP 5.37 | GRP1.8 | Exp 9 | 93.3 | a | 1750.0 | a | 0.0 | 93.3 | a |
| GRP 5.3 | GRP1.8 | Exp 9 | 93.3 | a | 1913.3 | a | 0.0 | 93.3 | a |
| GRP 8.15 | GRP1.8 | Exp 9 | 86.7 | a | 1970.0 | a | 0.0 | 86.7 | a |
| GRP 7.18 | GRP1.8 | Exp 9 | 93.3 | a | 2076.7 | a | 0.0 | 93.3 | a |
| II 10.29 | 35S | Exp 10 | 40.0 | **b** | 503.3 | **c** | 13.3 | 53.3 | **b** |
| Cruza 148 | control | Exp 10 | 53.33 | **b** | 793.3 | **c** | 0.0 | 53.3 | **b** |
| II 4.23 | 35S | Exp 10 | 73.3 | **b** | 870.0 | **c** | 13.3 | 86.7 | a |
| II 8.5 | 35S | Exp 10 | 60.0 | **b** | 930.0 | **c** | 20.0 | 80.0 | a |
| II 12.3 | 35S | Exp 10 | 80.0 | a | 1100.0 | **b** | 0.0 | 80.0 | a |
| II 6.45 | 35S | Exp 10 | 100.0 | a | 1143.3 | **b** | | 100.0 | a |
| II 6.11 | 35S | Exp 10 | 86.7 | a | 1320.0 | **b** | 13.3 | 100.0 | a |
| II 2.10 | 35S | Exp 10 | 86.7 | a | 1403.3 | **b** | 0.0 | 86.7 | a |
| II 5.8 | 35S | Exp 10 | 73.3 | **b** | 1440.0 | **b** | 0.0 | 73.3 | **b** |
| II 4.32 | 35S | Exp 10 | 86.7 | a | 1530.0 | a | 6.7 | 93.3 | a |
| II 11.2 | 35S | Exp 10 | 93.3 | a | 1590.0 | a | 0.0 | 93.3 | a |
| II 11.5 | 35S | Exp 10 | 86.7 | a | 1613.3 | a | 6.7 | 93.3 | a |
| II 3.32 | 35S | Exp 10 | 93.3 | a | 1763.3 | a | 0.0 | 93.3 | a |
| II 4.5 | 35S | Exp 10 | 86.7 | a | 1813.3 | a | 0.0 | 86.7 | a |
| II 11.22 | 35S | Exp 10 | 93.3 | a | 1856.7 | a | 0.0 | 93.3 | a |
| II 14.4 | 35S | Exp 10 | 93.3 | a | 1903.3 | a | 0.0 | 93.3 | a |
| II 13.2 | 35S | Exp 10 | 93.3 | a | 1953.3 | a | 0.0 | 93.3 | a |
| II 5.22 | 35S | Exp 10 | 93.3 | a | 2030.0 | a | 0.0 | 93.3 | a |

| | | | | | | | | | | | | |
|---|---|---|---|---|---|---|---|---|---|---|---|---|
| Desiree | control | Exp 10 | 93.3 | | a | 2056.7 | | a | 0.0 | 93.3 | | a |
| II 6.48 | 35S | Exp 10 | 93.3 | | a | 2066.7 | | a | 0.0 | 93.3 | | a |
| II 10.24 | 35S | Exp 10 | 100.0 | | a | 2173.3 | | a | | 100.0 | | a |
| II 3.28 | 35S | Exp 10 | 100.0 | | a | 2333.3 | | a | | 100.0 | | a |
| Cruza 148 | control | Exp 11 | 6.7 | 6.5 | **b** | 943.3 | 6.0 | **c** | 0.0 | 6.7 | 4.2 | **c** |
| II 10.31 | 35S | Exp 11 | 6.7 | 6.5 | **b** | 1046.7 | 7.2 | **bc** | 13.3 | 20.0 | 7.5 | **bc** |
| Desiree | control | Exp 11 | 33.3 | 15.5 | ab | 686.7 | 14.5 | abc | 6.7 | 40.0 | 17.0 | abc |
| GRP 6.58 | GRP1.8 | Exp 11 | 33.3 | 16.7 | ab | 620.0 | 14.8 | abc | 6.7 | 40.0 | 17.2 | abc |
| GRP 8.10 | GRP1.8 | Exp 11 | 33.3 | 16.7 | ab | 700.0 | 15.5 | abc | 20.0 | 53.3 | 22.0 | ab |
| II 10.18 | 35S | Exp 11 | 33.3 | 16.7 | ab | 126.7 | 17.0 | abc | 0.0 | 33.3 | 15.5 | abc |
| II 4.44 | 35S | Exp 11 | 46.7 | 21.5 | ab | 493.3 | 21.2 | abc | 0.0 | 46.7 | 20.5 | ab |
| II 5.11 | 35S | Exp 11 | 53.3 | 24.3 | a | 423.3 | 23.0 | ab | 0.0 | 53.3 | 23.7 | a |
| II 11.3 | 35S | Exp 11 | 46.7 | 21.8 | ab | 400.0 | 23.5 | a | 0.0 | 46.7 | 20.5 | ab |
| II 6.29 | 35S | Exp 11 | 60.0 | 23.7 | a | 933.3 | 25.7 | a | 0.0 | 60.0 | 23.0 | ab |
| GRP 6.54 | GRP1.8 | Exp 11 | 53.3 | 22.8 | a | 436.7 | 25.8 | a | 0.0 | 53.3 | 22.0 | ab |
| II 5.19 | 35S | Exp 11 | 80.0 | 29.3 | a | 33.3 | 27.8 | a | 0.0 | 80.0 | 29.0 | a |
| II 4.16 | 35S | Exp 12 | 33.3 | 10.2 | ab | 430.0 | 6.5 | **b** | 13.3 | 46.7 | 12.7 | ab |
| CRUZA 148 | control | Exp 12 | 26.7 | 6.8 | **b** | 440.0 | 7.2 | **b** | 6.7 | **33.3** | 7.7 | **b** |
| II 6.3 | 35S | Exp 12 | 33.3 | 8.7 | **b** | 670.0 | 9.5 | **b** | 13.3 | 46.7 | 12.0 | ab |
| II 10.29 | 35S | Exp 12 | 33.3 | 10.2 | ab | 670.0 | 11.7 | ab | 0.0 | **33.3** | 7.7 | **b** |
| GRP 7.15 | GRP1.8 | Exp 12 | 52.8 | 12.8 | ab | 1145.8 | 14.5 | ab | 0.0 | 52.8 | 10.8 | ab |
| GRP 3.11 | GRP1.8 | Exp 12 | 46.7 | 15.3 | ab | 856.7 | 14.7 | ab | 6.7 | 53.3 | 14.8 | ab |
| GRP 3.45 | GRP1.8 | Exp 12 | 57.8 | 15.0 | ab | 1105.6 | 15.0 | ab | 0.0 | 57.8 | 13.0 | ab |
| Desiree | control | Exp 12 | 80.0 | 21.0 | a | 1760.0 | 21.0 | a | 6.7 | 86.7 | 21.3 | a |

* Area under disease progress curve

** Transgenic events with 100% wilting were not tested for latent infection
